# The reliability of acoustic classification models for determining avian vocalisation patterns

**DOI:** 10.64898/2025.12.15.694294

**Authors:** O.C. Metcalf, C. Alencar Nunes, W.A. Hopping, A.C. Lees, V. Lostanlen, J. Barlow

**Affiliations:** Lancaster Environment Centre, Lancaster University, LA1 3FQ; Department of Natural Sciences, Manchester Metropolitan University, M1 5GD, UK; Cornell Lab of Ornithology, Cornell University, Ithaca, NY 14850, USA; LS2N - Laboratoire des Sciences du Numérique de Nantes; LS2N - équipe SIMS - Signal, IMage et Son

**Keywords:** ecoacoustics, ornithology, passive acoustic monitoring, machine learning

## Abstract

Automated detection and classification of species vocalisations offers the potential to utilise acoustic datasets across unprecedented spatial and temporal scales. However, classification algorithms inevitably generate errors, and error rates vary with context. While methods for quantifying error rates in ecoacoustics are well established, there is limited research on what level of model performance is sufficient to reliably determine avian vocalisation patterns. Using an extensive fully expert-labelled acoustic dataset from Peru (18 hours, 6 sites), we examined changes in the probability of detecting target bird vocalisations in the first hour after dawn to address three key questions: (1) How sensitive are models predicting detection probability over time to reductions in classification accuracy? (2) To what extent does aggregating detections over longer time periods impact classification accuracy? Additionally, we used the labelled dataset to assess how the creation and composition of a test dataset for assessing classifier performance can impact the reliability of accuracy metrics: (3) Are estimates of classification precision robust when test and deployment datasets are not independent and identically distributed?

Our results indicate that poor classification performance—especially low precision—can lead to misleading inferences about temporal patterns of vocalisations. Aggregating classifier predictions over longer time periods improved recall but often resulted in misleading patterns of vocal behaviour by reducing precision and temporal resolution. We also demonstrate that precision can be substantially overestimated when species presences are rarer in the deployment dataset than in test data. These findings highlight the importance of cautious application of automated classification in acoustic ecology and the need for accuracy assessment methods tailored to the intended ecological analysis.

## Introduction

Automated detection and classification of vocalisations of target species has been proposed as an effective method for analysing acoustic datasets at large spatiotemporal scales (Stowell and Sueur, 2020). The maturation of global, publicly available, and adaptable classification algorithms such as BirdNET (Kahl et al., 2021) and Google Perch (Denton et al., 2023) is bringing this reality closer. However, although these algorithms have already been hailed as effective at classifying species on a global scale (e.g. Bota et al., 2023. Funosas et al., 2024, Sethi et al. 2024), questions remain about performance in speciose regions (e.g. de Araújo, 2024). Accurate classification of presence or absence - whether or not a target sound occurs in a predetermined time period - over large datasets is key for successful use of the data in downstream modelling, allowing ecological inferences across a wide variety of questions, such as variation in species occurrence, relative abundance, or vocal activity over time (Brunk et al., 2025, Cole et al., 2022, Pérez-Granados, 2023, Sethi et al., 2024).

As statistical models, classification algorithms will generate both false positive and false negative errors (see the glossary in Table 1 for definitions of technical terms), with error rates varying by circumstance. Whilst there is a substantial body of literature looking at how to quantify error rate (Knight et al., 2017, Van Merriënboer et al., 2024, Wood and Kahl, 2024), there is only limited research assessing what level of model performance is good enough to reliably determine avian vocalisation patterns from classification predictions (Kitzes et al., 2025). Research on this topic has tended to focus around handling classification errors through estimating detectability using occupancy modelling (Balantic et al., 2019, Chambert et al., 2018, Ogawa et al., 2025, Rempel et al., 2019., Wright et al., 2020), and has found that this approach can be robust to some false positives and false negative errors, but with false positives generally considered more problematic (Wood and Kahl 2024).

While the attention paid to undesirable false positives is important (Kershenbaum et al., 2025), much less attention has been devoted to the rate of false negatives, otherwise known as recall (Table 1). This is important as reductions in recall have the potential to influence ecological inferences by underestimating species presence, or if false negatives are unevenly distributed, biasing detection histories. Additionally, little research has been done on how classification errors impact other forms of ecological modelling that do not include direct estimations of detectability, such as the assessment of changing vocal behaviour or non-vocal sounds over time.

Here, we present a case study using a fully expert labelled acoustic dataset from Peru (Hopping et al., 2022) analysed to assess changes in the probability of detecting target bird species vocalisations in the first hour from dawn. Understanding temporal changes in vocalisation behaviour is highly valuable in ecological research - both as an important aspect of natural history and animal behaviour, but also as an important component in assessing species detectability and for optimising survey protocols. We use these temporal questions to elucidate three challenges related to the assessment of classification error determining avian vocalisation patterns from ecoacoustic datasets:

### Question 1: How sensitive are patterns of avian vocalisations derived from automated classification models to reductions in precision and recall?

When using automated classification data to model changes in vocalisation behaviour, it is necessary to determine a threshold for classification accuracy prior to accepting the classification data for further ecological modelling. This involves a necessary trade off between precision and recall (Table 1). While researchers may be tempted to maximise precision to avoid false positives, the changes in recall could also have an important effect on ecological patterns. We quantify the level of classification error that still permits robust ecological modelling of temporal activity patterns, and assess whether reducing precision or recall has a greater negative impact on the reliability of avian vocalisation patterns over the course of the dawn chorus.

### Question 2: To what extent can summarising detection histories over longer time periods mitigate poor recall?

One of the reasons occupancy models can be less sensitive to low recall is that only a single presence is required over a set period - so if a recording contains multiple target sounds during that period, the classification algorithm only needs to detect one of them reliably. This process can be replicated in temporal or spatial analysis by summarising species presence over multiple classification windows. For instance, if a species vocalises 10 times per minute, a recall of 0.1 should still maintain an accurate representation of the species’ vocal presence at the 1 minute scale. We therefore evaluate whether aggregating temporal samples can mitigate the effects of low recall, and improve models of temporal activity patterns.

### Question 3: Are precision estimates robust to violations of the assumption that the distribution of presence to absence cases are identically distributed data between test and deployment datasets?

Calculating precision accurately requires that the ratio of presence to absence be similarly distributed between test and deployment datasets. Test datasets also need to contain enough positive and negative cases to robustly generate the required accuracy metrics, and yet be sufficiently small that the entire dataset can be manually labelled. The best way to ensure a representative distribution of presence to absence in the test dataset is to sample randomly from the deployment dataset - yet when target sounds are rare, a large sample size is needed to obtain enough positive cases to rigorously assess precision. Manually labelling such large test datasets is highly time consuming, and can be a major constraint on passive acoustic monitoring projects. Strategies to increase the number of presences and reduce the total test dataset size can therefore be tempting - for instance drawing test data from the peak of acoustic activity (i.e. the dawn chorus for most birds), or from a subset of recorders where the species is known to be present. However, such strategies can cause distribution shifts between the test and distribution datasets. Whilst the impact on precision is a mathematical property of the change in distribution between the two datasets, we use nine example species from our Peru dataset to illustrate how estimated and achieved precision shift with differences in distribution between the test and deployment dataset, even while classification performance remains the same.

## Methods

### Box 1. Assessing the performance of classification algorithms: key methods, challenges and a glossary of terms

Classification algorithms in ecoacoustics provide predictions of the presence or absence of a target sound over a set time period in the form of scores, often referred to as confidence scores (typically 0-1). However, these scores are not probabilities, and are not comparable between species or even across studies of the same species (Wood and Kahl, 2024). This means that classification performance should be assessed in each ecoacoustic study, even when using pre-existing classification models such as BirdNET. There are a number of approaches to assessing classification performance, with most converting classification scores into binary presence and absence classes using a threshold, before generating a range of performance metrics.

Selection of performance metrics in ecoacoustics is complicated due to class imbalance (Knight et al., 2017), with absences typically more common than presences. Passive acoustic studies are often conducted at large spatiotemporal scales - in which target species will not be present at every survey site, and recording schedules may include periods when target species do not vocalise. Moreover many species are more often silent than vocalising even during peak communication periods. In addition, the optimal length of audio for classification algorithms to detect species presence tends to be short; BirdNET uses 3s windows as an optimal period for accurate classification of most bird species, potentially resulting in a high number of windows in which targets are absent.

Many metrics used to assess classification accuracy, such as balanced accuracy, sensitivity and specificity, assume that i) both positive and negative classes are equally common in the dataset and ii) both types of correct classification are equally valuable and detrimental (Tharwat, 2018). Class imbalance in ecoacoustic datasets means that these metrics are either less informative or entirely inappropriate for imbalanced datasets (de la Cruz Huayanay et al., 2023) such as those in ecoacoustics. This is generally recognised in the field, and papers offering guidance on assessing classification performance (e.g. Knight et al., 2017) have recommended the use of appropriate metrics for imbalanced data. In particular, Precision, Recall, and metrics derived from these such as F score and Area Under the Precision-Recall Curve, and are now widely adopted in ecoacoustics to assess classification performance (Perez-Granados, 2023).

[End Box 1]

**Table 1.**
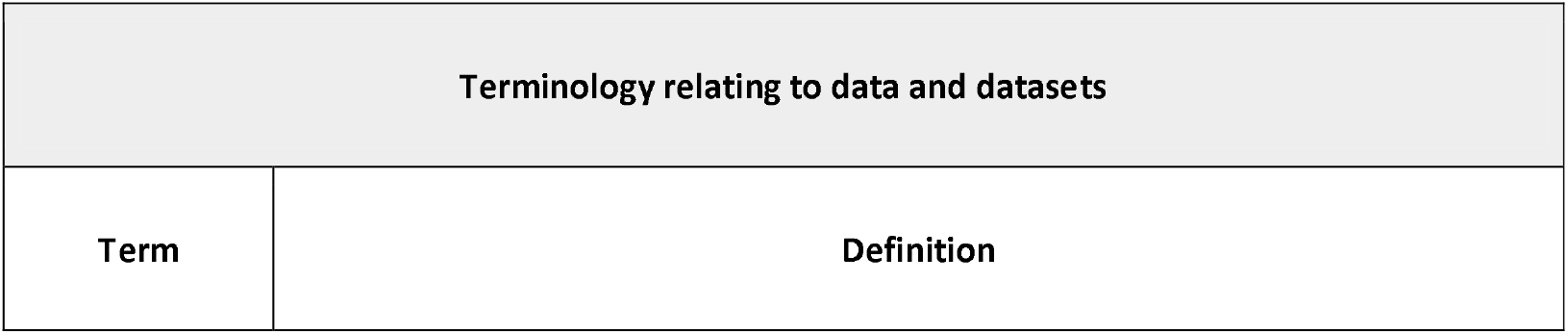
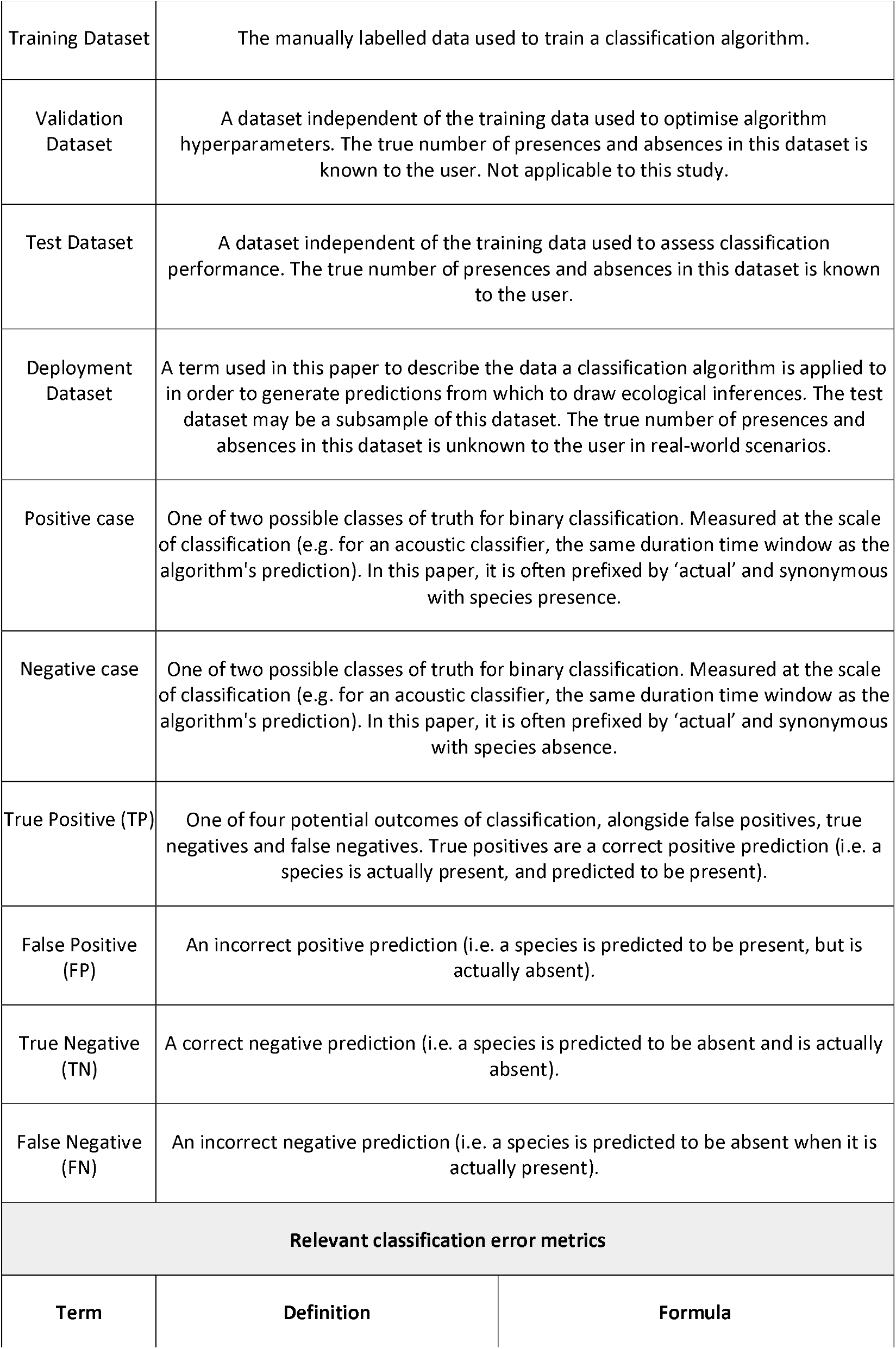
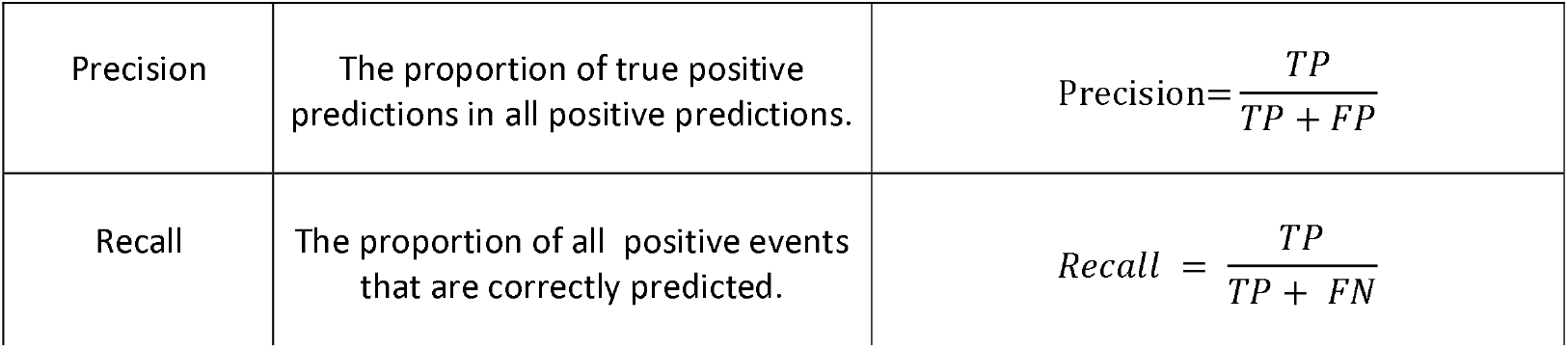
A glossary of key terms relating to classification accuracy.

| Terminology relating to data and datasets |  |
| --- | --- |
| Term | Definition |

|  |  |  |
| --- | --- | --- |
| Training Dataset | The manually labelled data used to train a classification algorithm. |  |
| Validation Dataset | A dataset independent of the training data used to optimise algorithm hyperparameters. The true number of presences and absences in this dataset is known to the user. Not applicable to this study. |  |
| Test Dataset | A dataset independent of the training data used to assess classification performance. The true number of presences and absences in this dataset is known to the user. |  |
| Deployment Dataset | A term used in this paper to describe the data a classification algorithm is applied to in order to generate predictions from which to draw ecological inferences. The test dataset may be a subsample of this dataset. The true number of presences and absences in this dataset is unknown to the user in real-world scenarios. |  |
| Positive case | One of two possible classes of truth for binary classification. Measured at the scale of classification (e.g. for an acoustic classifier, the same duration time window as the algorithm's prediction). In this paper, it is often prefixed by 'actual' and synonymous with species presence. |  |
| Negative case | One of two possible classes of truth for binary classification. Measured at the scale of classification (e.g. for an acoustic classifier, the same duration time window as the algorithm's prediction). In this paper, it is often prefixed by 'actual' and synonymous with species absence. |  |
| True Positive (TP) | One of four potential outcomes of classification, alongside false positives, true negatives and false negatives. True positives are a correct positive prediction (i.e. a species is actually present, and predicted to be present). |  |
| False Positive (FP) | An incorrect positive prediction (i.e. a species is predicted to be present, but is actually absent). |  |
| True Negative (TN) | A correct negative prediction (i.e. a species is predicted to be absent and is actually absent). |  |
| False Negative (FN) | An incorrect negative prediction (i.e. a species is predicted to be absent when it is actually present). |  |
| Relevant classification error metrics |  |  |
| Term | Definition | Formula |
| Precision | The proportion of true positive predictions in all positive predictions. | $\text{Precision} = \frac{TP}{TP + FP}$ |
| Recall | The proportion of all positive events that are correctly predicted. | $\text{Recall} = \frac{TP}{TP + FN}$ |

To assess how avian vocalisation patterns are impacted by classification error, we assessed when target tropical bird species are most likely to vocalise in the first hour after sunrise using a fully labelled dataset. We used the publicly available dataset (Hopping et al., 2022) from the Inkaterra Reserva Amazonica, Peru (12°32’07.8”S, 69°02’58.2”W, hereafter the Case Study dataset). The dataset comprises 18 1-hour recordings from 05:00-06:00 PET, collected simultaneously at six sites on three days between January 14 and February 2, 2019. All bird vocalisations in the recordings were identified where possible and labelled by WAH (see Hopping et al., 2022 for full details). 131 bird taxa were detected in the dataset, of which 93 species were identified to species level and detected more than ten times, and therefore included in the dataset for this study.

The presence labels for these species were converted into detection histories for every 3s period using an adapted version of the code provided by Hopping et al., (2023). Three seconds is the same period that BirdNET provides confidence scores for, and is the unit used to measure species presence/absence throughout. Each species therefore has 18 presence/absence histories, with each history having a length of 1200 3s time windows. Note that this should be considered realistic examples of ecological truth but not an actual study of these species behaviour - for instance we have made no attempt to model spatial or temporal autocorrelation in the data, and we are treating 05:00 as synonymous with sunrise for the purposes of these simulations. For an in depth analysis of temporal variation in avian vocalisation using this dataset see Hopping et al., (2023).

### Question 1: How sensitive are patterns of avian vocalisations derived from automated classification models to reductions in precision and recall?

To assess the sensitivity of avian vocalisation patterns to classification errors in the modelled data, we modelled the vocalisation history of a representative subset of species, adding false positives and false negatives to the vocalisation history to simulate different levels of classification error. All analysis was conducted in *R* (v4.5.1, R Core Team, 2025).

For Questions 1 and 2, we analysed the same subset of species. Species were selected by first restricting the case study dataset to those with an average detection rate exceeding 20 presences per hour. Next we used two variables to select the species - vocalisation abundance (i.e. presence to absence ratio of 3s windows) and a combined measure of the propensity to vocalise in series or duration of vocalisation (i.e. the average number of consecutive presences once a bird starts vocalising), as we believe these variables may interact with recall rates to impact vocalisation probability over time. We selected species that allowed a representation of each combination of low, medium, and high values for each variable. To do so, we created subsets of the species pool for taxa that had vocalisation abundances lower than the 1st quartile, between the 1st and 3rd quartile, and greater than the first quartile. Then for each subset, three species were selected with low, medium and high values of vocalisation clustering. Species were subjectively chosen as representing the best combinations of vocalisation abundance and vocalisation clustering, and to provide phylogenetic diversity.

**Table 1.**
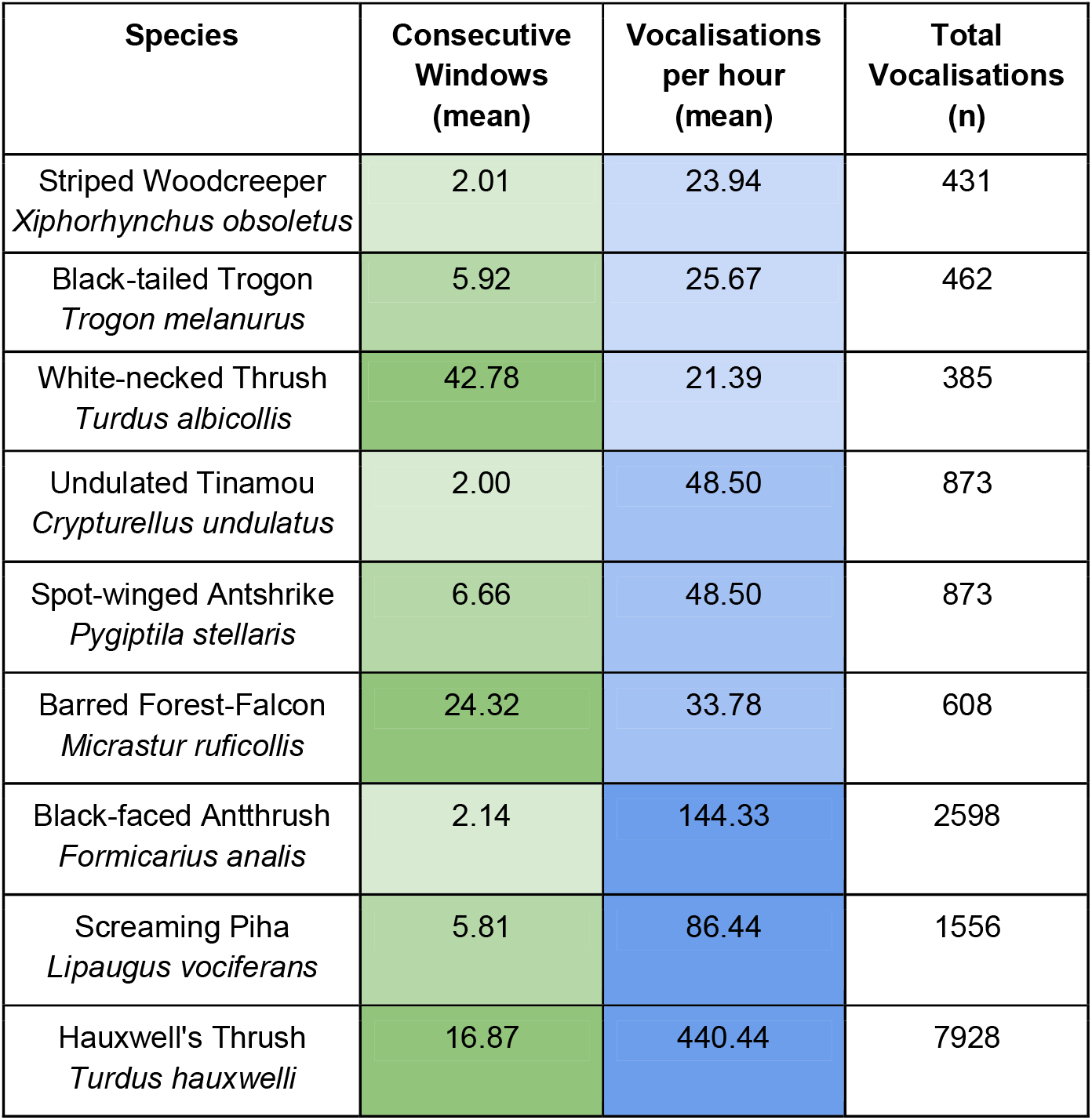
Target species for Questions 1 and 2, with measures of propensity for vocalising in series and/or vocalisation duration, vocalisation abundance, and total vocalisations labelled in the Case Study dataset. Species order is grouped by mean vocalisations per hour, then ordered within each group by mean consecutive windows.

To simulate classification error, we used predetermined precision and recall scores of 0.1, 0.25, 0.5, 0.75, 0.9 and 1. We tested all combinations of these scores, calculating the required number of false positives and negatives required for each detection history to obtain the required precision and recall scores. We assigned false positives by changing true negatives at randomly selected time slots to positives, and false negatives by changing true positives to negatives. Models with both precision and recall scores of 1 had no classification error, no false positives or negatives were introduced to the detection history, and are treated as the reference data against which the impact of classification error can be assessed. We repeated the simulation of classification error ten times for each species and pairwise level of error, so that true positives and negatives in different time windows were altered in each repetition. For some species simulating very low levels of precision was not possible as there were not enough true negatives to change to false positives to achieve the desired precision scores. These were 0.1 precision for *Formicarius analis* and 0.1 and 0.25 precision for *Turdus hauxwelli*.

To assess the impact of classification error on avian vocalisation patterns, we independently modeled the probability of detection in each time window for every species, using the simulated detection histories (glmmTMB package v1.1.9, Brooks et al., 2017). We modeled the observed presence (1) or absence (0) of the species (Y) as a function of time, its quadratic term, precision, and recall, including all interaction terms: Y ∼ (time + I(time^2^)) × precision × recall + (1 | repetition). The model was fitted using a binomial error distribution with a logit link. Time since dawn was included as a continuous variable, defined as the end of each 3 second window, scaled to proportion of the hour (i.e., values from 0 to 1 in steps of 3/3600 ≈ 0.00083). We generated marginal predictions of detection probability over all values of time (ggeffects package v1.5.2, Lüdecke, 2018), and for all values of precision and recall.

We assessed the impact of precision and recall on our models by comparing all models with imperfect classification accuracy to a reference condition - the trends for each species using detection histories with perfect accuracy (e.g. precision = 1 and recall = 1, so that they are identical to the detection histories from the Case Study). Models of detection probability over time were considered affected by classification error if their linear or quadratic time trends differed significantly (p < 0.05) from the corresponding reference model. Marginal trends were estimated with emtrends (emmeans package v1.8.9; Lenth et al., 2024) using precision and recall as interacting factors, and compared to reference models via contrast. We then assessed whether reductions in precision or recall had greater influence by examining effect sizes and p-values from the quadratic time trend analysis, with effects deemed significant when p < 0.05.

### Question 2: To what extent can summarising detection histories over longer time periods mitigate poor recall?

To test the effect of summarising classification-derived detection histories over longer time periods on our phenology models, we summarised all of the simulated detection histories (presence-absences) at 1 minute and 5 minute scales. To summarise the 3s detection windows to a larger timescale, if a bird was present in any of the 3s windows within the longer summary window, the bird was assigned as present in the summary window. Only if a bird was absent in all of the 3s windows within a summary window was it assigned as absent.

Firstly, to see if summarising detections over longer periods resulted in improved recall without loss of precision, we recalculated precision and recall metrics at the new aggregated timescales of one and five minutes. We compared the mean and standard deviation of the precision and recall scores across all 10 repetitions from all three timescales. Secondly, we assessed whether aggregating over larger timescales reduces the impact of classification error on predicted detection probabilities. To do so, we repeated the analysis from Question 1 using the new aggregated detection histories. To avoid introducing timescale as an additional prediction variable that would necessitate a four-way interaction and overly complex models, we fitted the same binomial regression models as above for each combination of species and timescale aggregation. We used these models to predict marginal detection probabilities over the hour from dawn, plotting the trends from each time aggregation allowing for a visual comparison.

### Question 3: Are precision estimates robust to violations of the assumption that the distribution of presence to absence cases are identically distributed data between test and deployment datasets?

To examine the sensitivity of precision estimates to shifts in the prevalence of classes between test and deployment datasets, we defined four classification scenarios with presence to absence ratios of 2:1, 1:2, 1:100, and 1:1000 (e.g., the 2:1 scenario included 6,667 presences and 3,333 absences). Each scenario used a modest test dataset of 10,000 three-second predictions (8 h 20 min of audio), though the total number of detections had minimal impact. Classification performance was fixed across scenarios, with both precision and recall set at 0.9.

Next we projected how changes in the presence to absence ratios between the test and deployment dataset would cause precision to vary between that estimated from the test dataset and that achieved on the deployment dataset. To do so, we calculated the false positive rates (FPR) of the algorithm over the four test dataset scenarios as: FPR = (1- Precision) * Recall / (Precision * Ratio). We then projected each of these false positive rates on to deployment datasets with a range of presence to absence ratios varying from 1:1 to 1:2000, with the denominator increasing sequentially by 1. To maintain a realistic scenario for an ecoacoustics study, each deployment dataset was much larger than the test datasets, with detection histories of length n= 8,640,000 — equivalent to ten recorders deployed for 20 days and recording for 12 hours per day, broken into 3s units (as per BirdNET classifications). Combined with four test dataset scenarios, this gives a total of 8,000 classification scenarios, each with a unique combination of test dataset presence to absence ratio and deployment dataset presence to absence ratio.

Using the projected false positive rates, we calculated the number of true and false positives for the deployment dataset in each scenario. True positives were calculated as presences multiplied by recall, and false positives as absences multiplied by the false positive rate for each deployment dataset scenario. We then calculated the precision achieved on the deployment dataset using the standard formula (Table 1). For each presence:absence ratio in the test datasets, we plotted these new precision values against the presence:absence ratio in the deployment dataset using loss curves and a span of 0.005 with the geom_smooth function from the ggplot2 package (Wickham et al., 2016).

To illustrate the effect of distribution shift with real species, we plotted points on each of these curves that represent the actual presence:absence ratios for species vocalisations from the Case Study dataset. To select appropriately representative species for this analysis, we first calculated the presence:absence ratio over 3 s periods for all species, then calculated the decile values of the presence:absence ratio distribution. We selected species with ratios closest to the lowest and highest values, plus the 2nd, 4th, 6th and 8th deciles, and calculated how the precision would change for these species between each test dataset ratio and their respective presence:absence ratios in the Case Study dataset.

**Table 2:** Selected species, the number of time windows without a detection per window with a detection (odds of non-detection), and the decile of that odds ratio relative to the entire species pool.

| Species | Odds Ratio | Decile |
| --- | --- | --- |
| Ferruginous Pygmy-Owl<br>*Glaucidium brasilianum* | 1963.64 | 0% |
| Blue-headed Parrot<br>*Pionus menstruus* | 502.33 | 20% |
| Cobalt-winged Parakeet<br>*Brotogeris cyanoptera* | 186.21 | 40% |
| Plain-winged Antshrike<br>*Thamnophilus schistaceus* | 73.22 | 60% |
| Amazonian Grosbeak<br>*Cyanoloxia rothschildii* | 25.84 | 80% |
| Hauxwell's Thrush<br>*Turdus hauxwelli* | 2.72 | 100% |

To assess the scale of the challenge in obtaining a suitable test dataset for rare species or sounds, we calculated how many hours of data would be required to obtain a test dataset that maintained each species presence:absence ratio from the Case Study dataset, and contained 100 presences. To do so, we divided 100 by the vocalisation rate per hour of each species, and rounded up to the nearest hour.

## Results

### Question 1: How sensitive are patterns of avian vocalisations derived from automated classification models to reductions in precision and recall?

Our example species exhibit a range of vocalisation responses as time progresses after dawn when no classification error is simulated in the detection histories (Figure 1, red lines). Both the linear and quadratic terms for time are a significant predictor for all species (Appendix 1 for model summary tables). Four species show a concentrated peak in vocalisations during the hour, with peak detection probability for *Turdus albicollis* towards the start of the hour, *Xiphorhynchus obscurus* and *Pygiptila stellaris* in the middle and *Trogon melanurus* towards the end. Only *Lipaugus vociferans* showed a consistently increasing detection probability throughout the hour. Overall, detection probability is quite low, below 0.1 at all times for all species, except for the three species with the highest vocalisation abundance, of which *Turdus hauxwelli* has the highest detection probability – approaching 0.4 for the third quarter of the hour.

**Figure 1.**
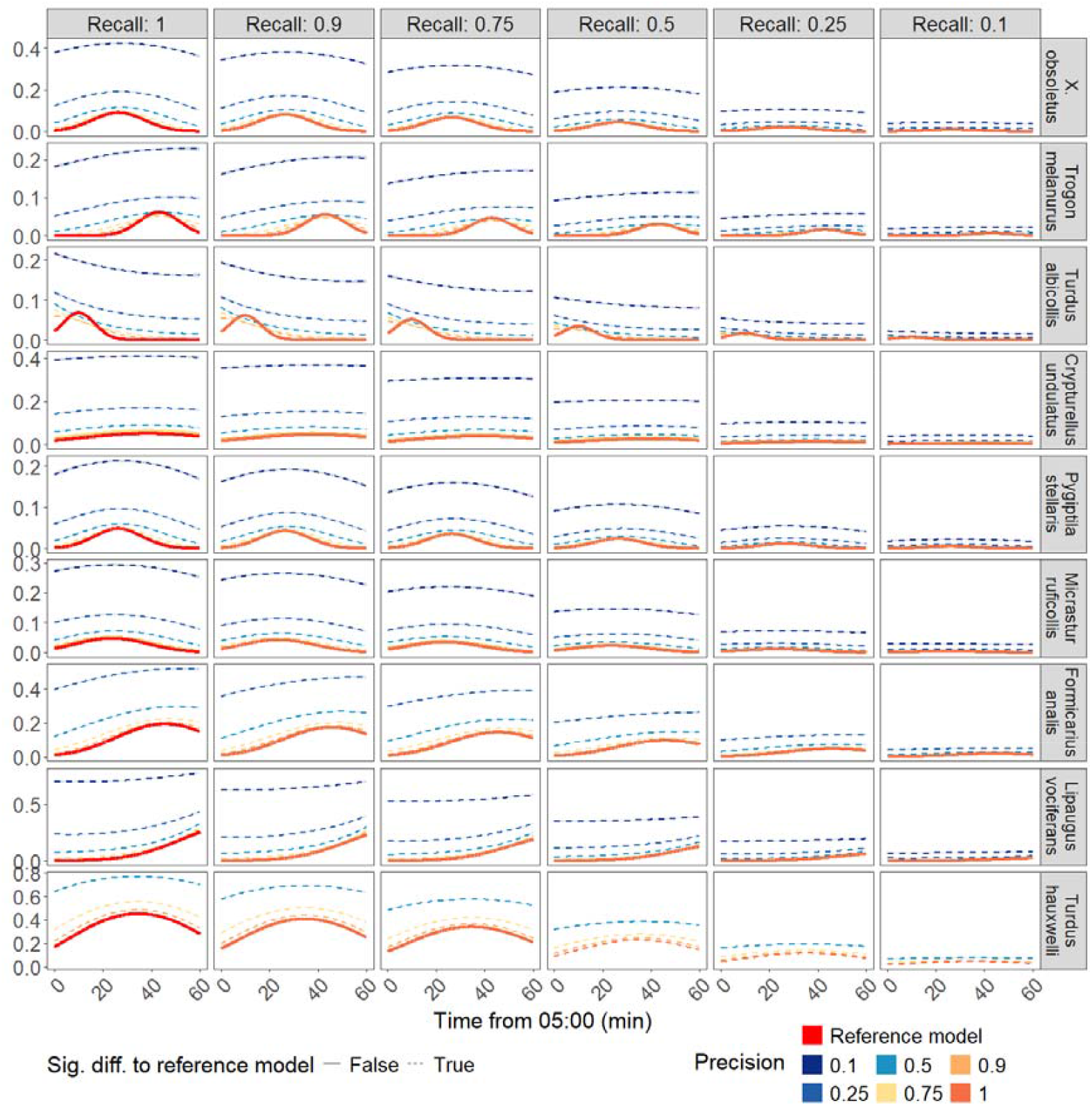
Modelled detection probability of the target species in the first hour from dawn. Reference conditions - trends with perfect accuracy (recall=1, precision=1), are shown in red. The recall score simulated in the modelled detection histories are shown in facet headers along the x axis, and precision values are illustrated with line colour. Trends that do not significantly differ from the reference value are shown as solid lines, dashed lines show trends that differ significantly from the trend for the reference condition.

Both imperfect precision and recall influenced the estimated relationship between time and detection probability (Figure 1), with most precision - recall combinations producing trends that differed significantly from the reference condition (*precision* = 1, *recall* = 1). When precision was perfect, most estimated trends (*n* = 51) did not differ significantly from the reference, except for *Turdus hauxwelli* with recall ≤ 0.5 (*n* = 3). The number of trends differing significantly from the reference increased sharply as precision declined: at a precision of 0.9, most trends (*n* = 42) were significantly different, except those for *Crypturellus undulatus* (all recall scores), *Micrastur ruficollis* (all recall scores except 0.9), and *Pygiptila stellaris* (recall = 0.1). In contrast, declining recall had a smaller effect, with 10 - 12 trends at each recall level remaining statistically indistinguishable from the reference condition.

Beyond statistical significance, when precision and recall scores exceeded 0.5, the trends displayed a high degree of visual similarity across all species, albeit with a reduced flattened trend lines at lower recall scores and increased detection probability with precision scores. This is especially true for the five species that did not show strong concentrated peaks in vocalisations. For the other four species, temporally concentrated patterns of vocalisations are lost as soon as precision declines below 1, even if recall is perfect. Similar reductions in recall preserve the temporal detail in detection probability, with distinct peaks in detectability still visually apparent even at recall scores as low as 0.5 for all four relevant species.

The post-hoc test shows that declining precision scores result in overestimated effect sizes, whether positive or negative. Declines in precision scores have a much greater impact on the estimated marginal effect size than similar declines in recall, and this disparity grows proportionately larger as classification scores reduce (Figure 2).

**Figure 2.**
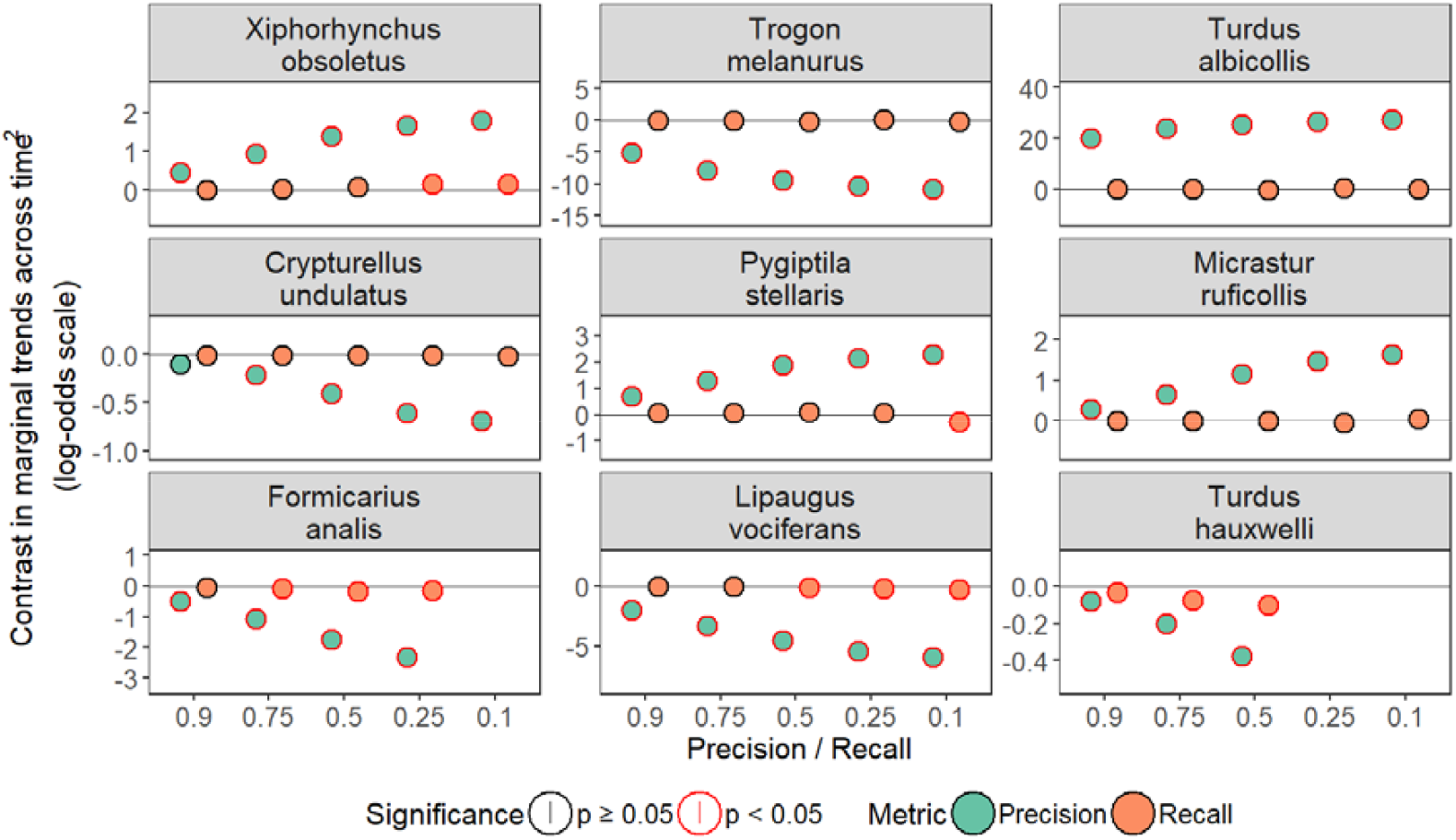
The estimated effect size of simulated recall and precision error levels on detection probability from dawn. Labels show p-values, with red text indicating significance at p<0.05.

### Question 2: To what extent can summarising detection histories over longer time periods mitigate poor recall?

Summarising detection histories over longer time periods substantially improves recall (Figure 3A), but decreases precision. Recall increases by a mean of 0.31 ± 0.23 when aggregating from 3 s to 1 min and 0.37 ± 0.29 when aggregating to 5 min, across all species and initial recall scores. In contrast, trends in precision are more variable (Figure 3B), differing by species and original recall score, but on average decreasing by −0.14 ± 0.17 between 3 s and 1 min and by −0.17 ± 0.25 between 3 s and 5 min. Precision responses also show evidence of an interaction between original precision and recall at the 3 s scale impacting the resultant precision at aggregated timescales. When original precision score is 0.1, precision generally increases with aggregation, except for *Xiphorhynchus obsoletus* and *Turdus albicollis* at 1 min, and *Turdus hauxwelli* at both 1 min and 5 min. As the original precision increases, changes in precision across timescales increasingly depend on the original recall: precision typically declines sharply when original recall is ≥ 0.75 (mean = −0.20 ± 0.19), declines more modestly when recall is ≤ 0.5 (mean = −0.08 ± 0.13), with some cases for *Trogon melanurus* even increasing. Of the 108 trends with an original precision value of 0.9 across all species, only 30 maintained a precision value above 0.75 when aggregated at longer time scales, and only 77 with a precision score above 0.5, reducing further to just seven and 43 respectively when only considering trends in which original recall values were also above 0.5.

**Figure 3.**
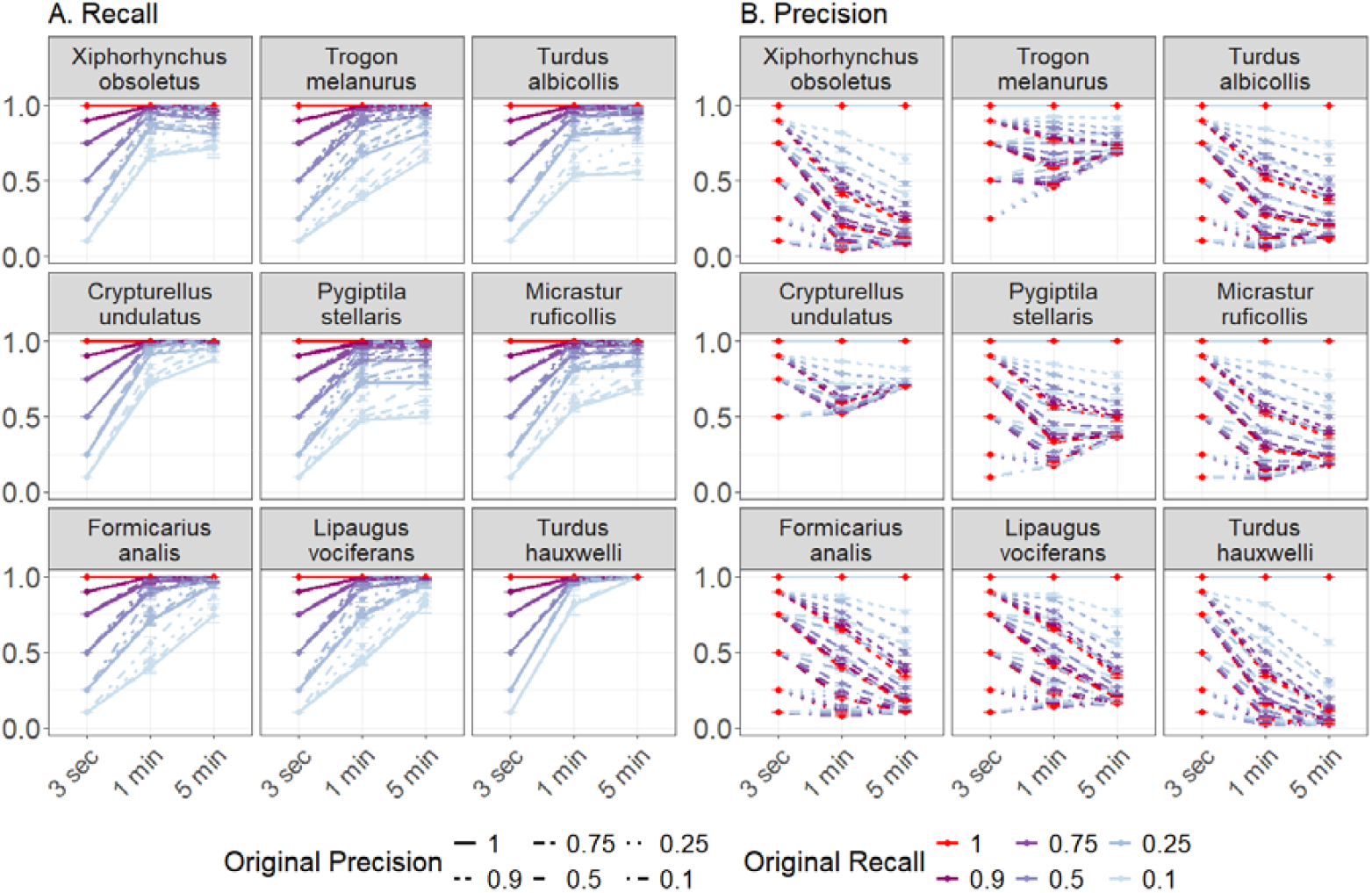
The effect on recall and precision scores of summarising data from 3 s time units (original recall values) to one minute and five minute units. The new recall and precision values are calculated taking mean values across all replicates of the simulated classification error, with error bars showing the standard deviation.

Aggregating detection histories over longer timescales substantially alters estimated temporal trends (Figure 4) - with decreases in classification accuracy resulting in increasingly divergent trends between timescales. Trends based on 3 s detection histories remain visually similar to the reference condition across all levels of classification accuracy. In contrast, trends for aggregated timescales increasingly deviate not only from the reference condition but also from their counterparts at perfect accuracy once precision and recall fall below 0.9. Notably, these differences are not solely attributable to classification error. Even at perfect accuracy, aggregating detection histories over longer timescales alters the estimated trends relative to the reference condition, particularly for the 5 min aggregation and for species with the highest vocalisation abundance—*Formicarius analis, Lipaugus vociferans*, and *Turdus hauxwelli* which are predicted as vocalising in all samples.

**Figure 4.**
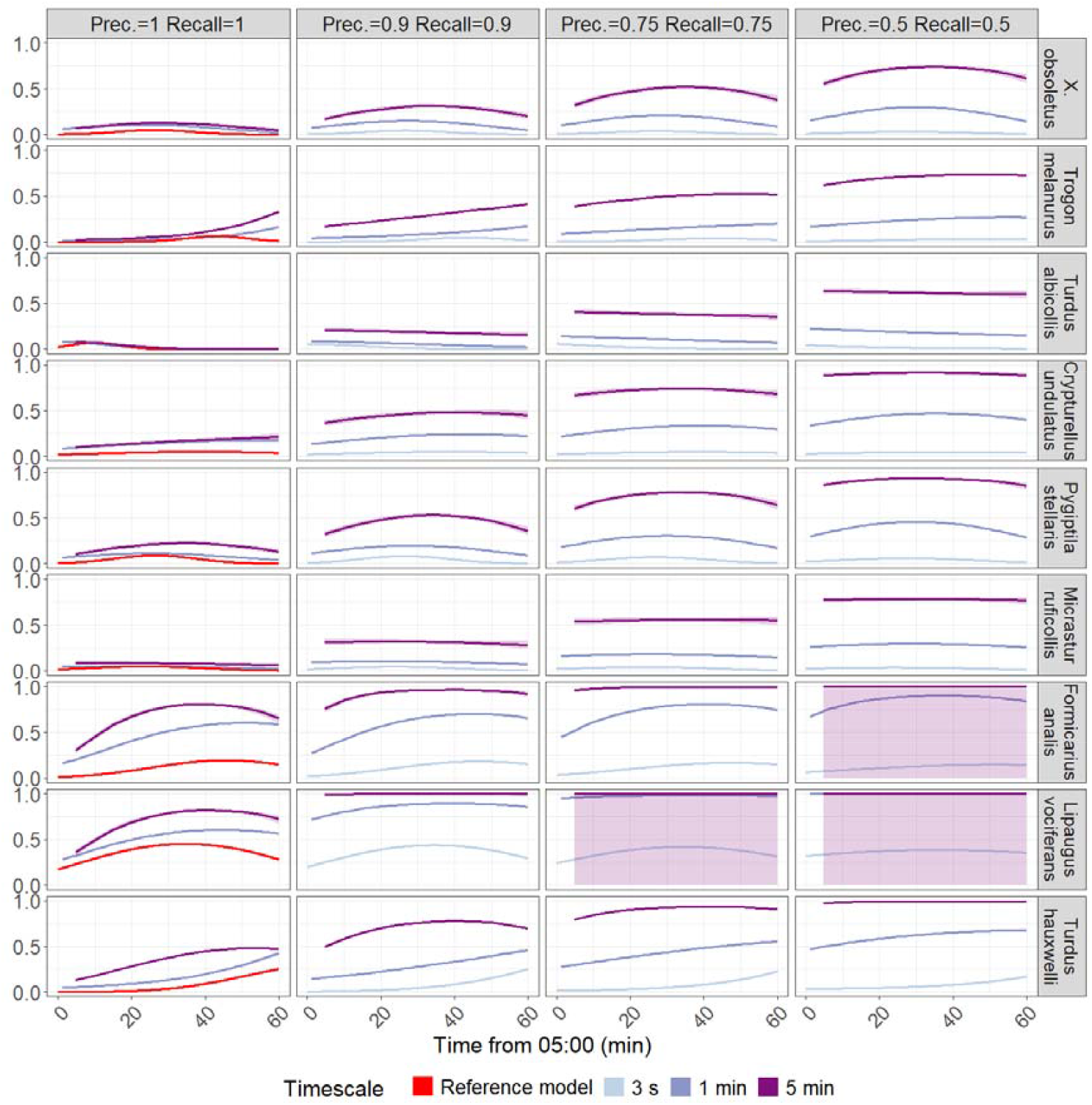
A comparison of predicted vocalisation probabilities when detections are aggregated at three timescales - the original 3 s prediction intervals (red when accuracy is perfect, otherwise light blue lines), 1 min (mid blue lines) and 5 minutes (purple lines) - and across four levels of decreasing classification accuracy (horizontal facets). Accuracy scores given here are those for the 3 s detection histories.

### Question 3: Are precision estimates robust to violations of the assumption that the distribution of presence to absence cases are identically distributed data between test and deployment datasets?

Precision declines dramatically when the ratio of presences to absences is much greater in the test dataset than in the deployment dataset. This is particularly apparent in the Test Datasets that have 2:1 and 1:2 presence/absence ratios. In the former case, all of our example species precision scores declined from 0.9 to below 0.1 except for *Turdus hauxwelli* and *Cynanoloxia rothschildii* (Figure 1A). In the latter case, most species retained precision scores above 0.1 except for *Brotogeris cyanoptera, Pionus menstruus*, and *Glaucidium brasilianum*, but all scores except *Turdus hauxwelli* declined by over 50%. For the example test sets with a 1:100 ratio, the three species with a vocalisation rate lower than 1:100 still showed declining precision, but as with the 1:1000 test set, all species with a lower ratio of presences to absences than the test set itself had precision scores overestimated by the test dataset. Figure 1B illustrates how much time is required to obtain an appropriately sized test dataset, containing 100 presences, for our example species. For four species the audio data required for manual assessment ranged between 1 and 16 hours, however *Pionus menstruus* and *Glaucidium brasilianum* required 42 and 164 hours of manually labelled data respectively.

## Discussion

The use of machine learning algorithms in ecoacoustics promises to allow the assessment of the presence of sonifying species on greater temporal and spatial scales than previously possible (Sethi et al., 2024), however in this study, we demonstrate that poor classification performance—particularly low precision—can lead to misleading avian vocalisation patterns. Moreover, we illustrate that precision can be easily overestimated when the statistical distribution of species presences differs between the test data and the real-world deployment context. Taken together, these findings highlight the need for caution when applying automated classification in acoustic analyses. While such models can offer substantial time savings (Manzano-Rubio et al., 2022, Cole et al., 2022), they may still demand considerable human oversight to rigorously evaluate performance and ensure the reliability of subsequent ecological interpretations of animal phenology.

Nevertheless, our assessment of temporal activity patterns exhibited a substantial degree of robustness to classification error. In most cases, models showed visually similar trends when precision scores were above 0.75, and 0.5 for some species, with recall scores at 0.5 or above. Although many of these trends were statistically significantly different from the reference models, the large detection history (n = 21,600) means that some differences that are detected as significantly different may not result in substantial differences in understanding ecological behaviour (e.g. Figure 1 for most species when recall and precision are greater than 0.5) - visual assessment of model similarity rather than over-reliance on p-values is warranted here. It is however worth bearing in mind that the random distribution of errors across the time-series in our simulations represent something of a best case scenario, as it mitigates against the introduction of bias. In real-world datasets, changes in environmental or ecological factors such as rainfall or co-occurrence of species with similar vocalisations, may affect accuracy (Kitzes et al., 2025) resulting in higher error rates at particular times of day and therefore biasing ecological models. Users of classification models should consider this risk and ensure that test datasets are suitably large to enable an evaluation of biases.

When compared to accuracy scores achieved in published studies using BirdNET, this suggests that classification accuracy will often, but not always, be suitable for obtaining accurate avian vocalisation patterns in phenological studies. A review by Pérez-Granados (2023) provides a good sample of studies using BirdNET on real-word datasets, albeit not necessarily for phenological studies. Of the nine studies included, with each covering multiple species, six had an average precision across all species estimated to be above 0.75. However, all the studies except one in North America, found that BirdNET precision scores were below 0.75 for at least one of the species studies, and below 0.5 for four of the studies. This suggests that ‘off-the-shelf’ classification models are capable of achieving precision scores suitable for modelling temporal activity patterns but further emphases the need for model performance to be assessed on a case-by-case basis.

Furthermore, only three of the nine studies assessed recall - of which two reported average recall below 0.5, and other studies have reported similar issues in obtaining high recall scores whilst maintaining high levels of precision (Funosas et al., 2024). While our analysis shows that lower recall has a smaller impact on ecological modelling than reduced precision, models with recall less than 0.5 consistently underestimated detection probability and exhibited flattened trends compared to models with perfect accuracy, even when precision was high. This suggests that studies only assessing precision but not recall (e.g.Sethi et al., 2024) may be inadvertently reporting trends that are significantly different to those that would be observed with better classification accuracy. It also underlines the importance of using methods to assess classification performance that are appropriate to the analysis being conducted.

Where classification models do not achieve high recall scores on a test dataset, aggregating data into longer time periods will lead to substantial improvement in recall. However, this comes with three important caveats. First, analysing time series at coarser resolutions than the outputs provided by classification models can significantly influence the interpretation of temporal patterns (Yoh et al., 2024). Second, our simulations show that increases in recall scores are often accompanied by simultaneous declines in precision. Even when precision and recall are high enough at initial timescales for avian vocalisation patterns to be robust to error, aggregation over longer timescales results in precision scores that suggest vocalisation patterns may no longer be reliable. Although the declines in precision are smaller than the increases in recall, the disproportionate impact of poor precision on phenological models mean that models from aggregated timescales differed drastically for most species from those at 3 s timescales. Third, manual assessment of classification performance at longer timescales is even more consuming, requiring reviewing longer periods of audio to achieve the same number of samples. In combination, these factors mean that we do not recommend summarising automated classification outputs over long time periods for phenological analysis, without further considerations in the process of binarizing classification prediction scores specific to timescale aggregation such as those recommended in Singer et al., (2025).

To effectively estimate accuracy metrics in phenological studies, it is necessary to derive them from a test dataset that is independent of, and with an identical distribution of classification classes to, the deployment dataset (Hand, 2012, Foody, 2023). We demonstrate that using test data that does not share the same distribution as the deployment dataset can lead to marked discrepancies between estimated and realised precision. Over-representation of positive cases can be tempting when trying to obtain a large enough sample size of vocalisations to satisfy the requirements of qualitative representation, as the addition of each positive case requires a disproportionately higher number of negative cases. Simple strategies for obtaining a high enough sample size, such as using test data from the dawn chorus when the deployment data spans the diel cycle can result in large divergences in distribution. The patterns in Figure 5A, while reflecting a simple mathematical relationship with changing class ratios, highlight a critical point: even modest shifts in the distribution of species presences can trigger steep increases in false positives (Foody et al., 2023), potentially undermining confidence in ecological conclusions drawn from automated classification.

**Figure 5:**
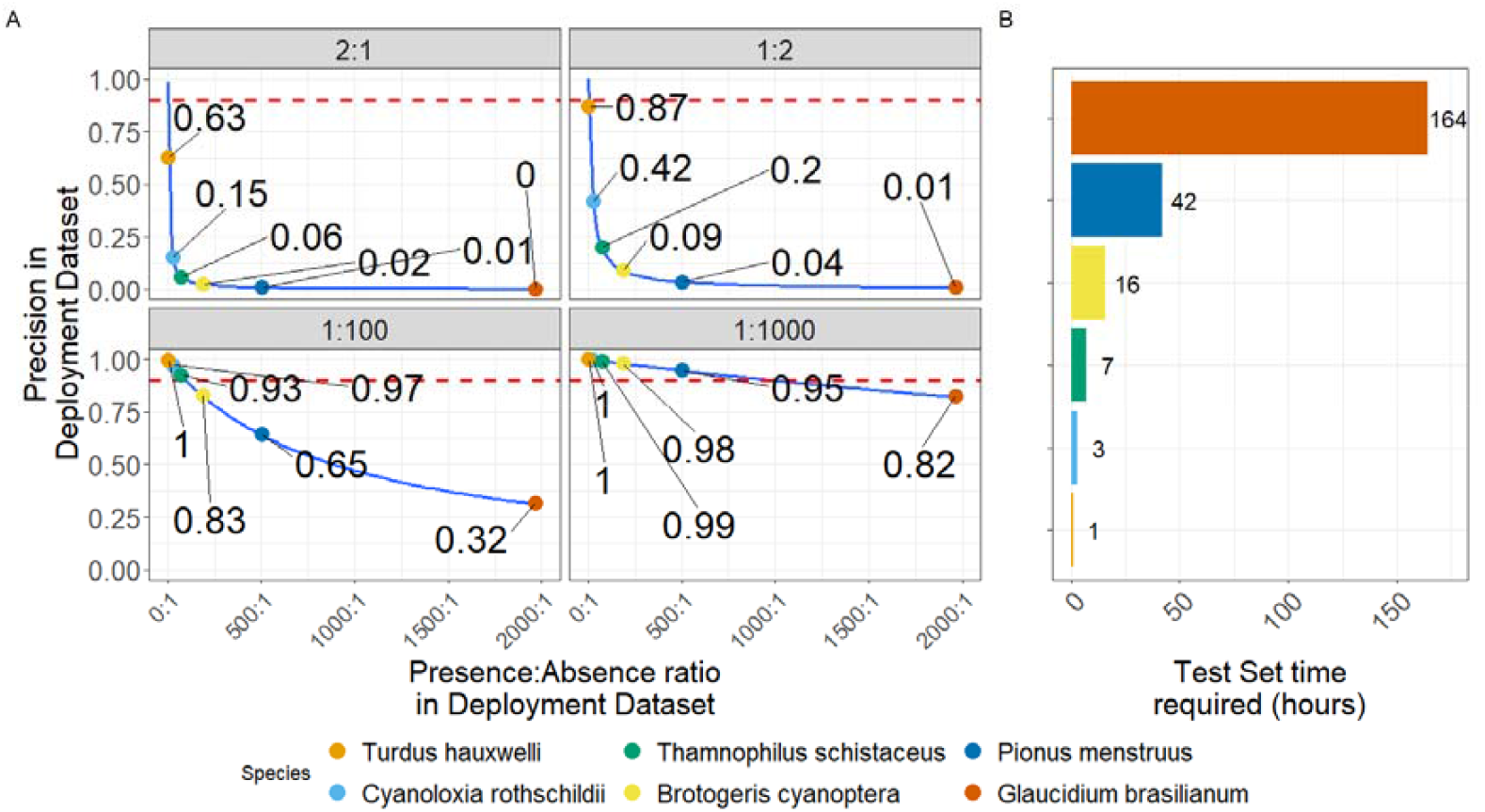
The impact of distribution shift between test and deployment datasets on precision values. **A:** Change in precision in the test dataset vs achieved in the deployment dataset Presence:absence odds ratio in the test dataset are shown in the facet headings, and for the deployment dataset on the x axis. Coloured points show how representative species from the Case Study dataset could be impacted. The red dashed line shows the assigned precision values for the Test Dataset of 0.9. **B**. The number of hours required for each species in a Test Dataset that both maintains the species vocalisation ratio in the Case Study dataset and contains 100 presences.

We show just how much audio data would be required to obtain a test dataset containing 100 vocalisations of each species, which vary by an order of magnitude depending on the species vocalisation abundance in the case study. This challenge poses two key problems for evaluating automated classification performance in passive acoustic monitoring. First, the vocalisation rate of a species—that is, how frequently it vocalises over a given period—is rarely known prior to data collection, and even when it is, vocalisation rates can vary substantially between habitats, across times of day and year, and in response to conspecifics, predators, or prey (Hutschenreiter et al., 2024). Such variability makes it difficult to determine in advance the sample size required for a robust test dataset. Second, for species with low vocalisation rates—whether due to rarity in the study area or limited vocal activity—the volume of audio that must be manually annotated to achieve sufficient sample size can be prohibitive. This is particularly limiting given that effective monitoring of rare species is often cited as a major advantage of passive acoustic monitoring (Klingbeil & Willig, 2015, Sugai et al., 2018). For example, in our case study, acquiring a test dataset containing 100 vocalisations of *Glaucidium brasiliensis* would require 164 hours of recordings. While this threshold of 100 vocalisations is arbitrary and performance could potentially be assessed with fewer vocalisations, determining the minimum sample size necessary remains an important area for further research. An additional consideration for multispecies or community studies, the test dataset size will be constrained by the species with the lowest vocalization rate, meaning studies may have to be limited to the most vocal species with subsequent biases, or invest in generating large test datasets.

As with the distribution of classification errors discussed in Questions 1 and 2, these test dataset size estimates represent a best-case scenario in which there is minimal covariate shift—that is, change in environmental or ecological conditions that affect both classification accuracy and vocalisation rates (Moreno-Torres et al., 2012, Metcalf et al., 2022) —across the deployment dataset. In practice, such stability is rare. Environmental conditions such as weather and background noise, as well as ecological factors including local abundance of conspecifics, predators, or prey, often vary across space and time. Where covariate shift is likely to alter error rates in ways that could confound ecological inference, test datasets may need to be drawn independently from multiple temporal or spatial subsets of the deployment. This substantially increases minimum sample size requirements. For instance, if classification accuracy for *Cyanoloxia rothschildii* needed to be evaluated separately for dawn and dusk at three locations, and vocalisation rates were constant, the required test dataset would grow from three hours to 18 hours of recordings. This is reflected in real-world uses of automated classification of acoustic data, where a study using automated classification models to assess breeding phenology in Marbled Murrelets *Brachyramphus marmoratus* required the manual assessment of 101,403 12 s clips (338 hrs) to establish classification performance at two distinct study areas (Duarte et al., 2024). In many cases, this additional manual annotation effort could be prohibitive. Accordingly, understanding how well automated classification models generalise, and identifying the covariates that drive changes in accuracy, should be considered a research priority in ecoacoustics to enable realistic assessment of the effort and cost required for such studies (Van Merriënboer et al, 2024). Valuable objectives for further research include gaining a fuller understanding of the level of manual labelling effort at which it becomes more efficient to use the manual labels for understanding patterns of vocalisation than continue with classification output. This would allow researchers to make informed decisions about whether automated classification of an entire dataset or manual review of a subsample is the better approach.

Where it is not possible to ensure that a test dataset is both a suitable size, independent and identically distributed to the deployment dataset, assessment of a sample of positive prediction cases as advocated by Wood and Kahl (2024) remains an effective alternative.

This approach allows the accurate and efficient assessment of precision, easy stratification of sampling across changing covariates, and removes the requirement for identical distribution as only positive cases are assessed. However, it is important to acknowledge that this approach to assessing classifier performance does not allow identification of false negatives, and therefore it is impossible to derive recall scores. Without this information, it is impossible to quantify the reliability of predicted absences and therefore negative predictions from data assessed in this manner should be treated as pseudo-absences. In turn this restricts the number of analysis methods suitable for this sort of data to those that directly assess detection probability in the modelling process, such as occupancy models (Kéry and Royle, 2020), and those that can use presence-only or presence/pseudo-absence data such as MaxEnt species distribution models (Renner and Warton, 2013).

## Conclusion

While automated classification offers powerful opportunities for large-scale acoustic monitoring, our findings emphasise that careful performance assessment is essential to avoid misleading patterns of vocalisation. Precision and recall thresholds for phenological modelling insensitive to classification error are often achievable but not guaranteed, and both covariate shift and inadequate or biased test datasets can severely compromise reliability. For rare or infrequently vocalising species, the effort required to obtain suitable test datasets may be prohibitive, underscoring the need for methods that balance accuracy, efficiency, and feasibility. Approaches such as targeted assessment of positive predictions can provide valuable precision estimates where ideal test datasets are impractical, but their limitations—particularly the inability to assess recall—must be acknowledged. Advancing ecoacoustics will require both methodological rigour in evaluating classification performance and a clearer understanding of what assessment methods are suitable for the desired analysis.

## Declaration of generative AI and AI-assisted technologies in the writing process

During the preparation of this work the author(s) used ChatGPT in order to improve the clarity of language. After using this tool/service, the author(s) reviewed and edited the content as needed and take(s) full responsibility for the content of the publication.

