## Supplementary materials for "The reliability of acoustic classification models for determining avian vocalisation patterns"

**Appendix 1: Model estimates by species**

| **Species** | **Term** | **Estimate** | **Std. Error** | **95% CI Lower** | **95% CI Upper** | **z value** | **p value** |
| --- | --- | --- | --- | --- | --- | --- | --- |
| Xiphorhynchus obsoletus | (Intercept) | -6.591 | 0.087 | -6.762 | -6.42 | -75.609 | <.001 |
| Xiphorhynchus obsoletus | I(time^2) | -18.819 | 0.438 | -19.676 | -17.961 | -42.996 | <.001 |
| Xiphorhynchus obsoletus | I(time^2):precision0.1 | 17.828 | 0.444 | 16.958 | 18.697 | 40.171 | <.001 |
| Xiphorhynchus obsoletus | I(time^2):precision0.1:recall0.1 | 3.098 | 1.656 | -0.148 | 6.345 | 1.871 | 0.061 |
| Xiphorhynchus obsoletus | I(time^2):precision0.1:recall0.25 | -0.913 | 0.961 | -2.797 | 0.971 | -0.95 | 0.342 |
| Xiphorhynchus obsoletus | I(time^2):precision0.1:recall0.5 | -0.976 | 0.745 | -2.436 | 0.484 | -1.31 | 0.19 |
| Xiphorhynchus obsoletus | I(time^2):precision0.1:recall0.75 | -0.547 | 0.668 | -1.857 | 0.763 | -0.818 | 0.413 |
| Xiphorhynchus obsoletus | I(time^2):precision0.1:recall0.9 | -0.3 | 0.64 | -1.555 | 0.955 | -0.469 | 0.639 |
| Xiphorhynchus obsoletus | I(time^2):precision0.25 | 16.213 | 0.452 | 15.327 | 17.099 | 35.856 | <.001 |
| Xiphorhynchus obsoletus | I(time^2):precision0.25:recall0.1 | 2.812 | 1.682 | -0.485 | 6.11 | 1.672 | 0.095 |
| Xiphorhynchus obsoletus | I(time^2):precision0.25:recall0.25 | -0.989 | 0.981 | -2.911 | 0.934 | -1.008 | 0.313 |
| Xiphorhynchus obsoletus | I(time^2):precision0.25:recall0.5 | -1.032 | 0.76 | -2.521 | 0.458 | -1.357 | 0.175 |
| Xiphorhynchus obsoletus | I(time^2):precision0.25:recall0.75 | -0.61 | 0.681 | -1.945 | 0.726 | -0.895 | 0.371 |
| Xiphorhynchus obsoletus | I(time^2):precision0.25:recall0.9 | -0.301 | 0.653 | -1.581 | 0.978 | -0.462 | 0.644 |
| Xiphorhynchus obsoletus | I(time^2):precision0.5 | 12.976 | 0.472 | 12.051 | 13.901 | 27.488 | <.001 |
| Xiphorhynchus obsoletus | I(time^2):precision0.5:recall0.1 | 3.252 | 1.741 | -0.16 | 6.664 | 1.868 | 0.062 |
| Xiphorhynchus obsoletus | I(time^2):precision0.5:recall0.25 | -1.114 | 1.028 | -3.128 | 0.9 | -1.084 | 0.278 |
| Xiphorhynchus obsoletus | I(time^2):precision0.5:recall0.5 | -0.996 | 0.795 | -2.555 | 0.563 | -1.253 | 0.21 |
| Xiphorhynchus obsoletus | I(time^2):precision0.5:recall0.75 | -0.424 | 0.712 | -1.819 | 0.971 | -0.595 | 0.552 |
| Xiphorhynchus obsoletus | I(time^2):precision0.5:recall0.9 | -0.2 | 0.682 | -1.536 | 1.136 | -0.294 | 0.769 |
| Xiphorhynchus obsoletus | I(time^2):precision0.75 | 8.4 | 0.51 | 7.401 | 9.4 | 16.471 | <.001 |
| Xiphorhynchus obsoletus | I(time^2):precision0.75:recall0.1 | 2.421 | 1.871 | -1.247 | 6.089 | 1.293 | 0.196 |
| Xiphorhynchus obsoletus | I(time^2):precision0.75:recall0.25 | -1.033 | 1.116 | -3.221 | 1.155 | -0.926 | 0.355 |
| Xiphorhynchus obsoletus | I(time^2):precision0.75:recall0.5 | -0.482 | 0.859 | -2.166 | 1.202 | -0.561 | 0.575 |
| Xiphorhynchus obsoletus | I(time^2):precision0.75:recall0.75 | -0.417 | 0.77 | -1.925 | 1.092 | -0.541 | 0.588 |
| Xiphorhynchus obsoletus | I(time^2):precision0.75:recall0.9 | -0.237 | 0.737 | -1.681 | 1.208 | -0.321 | 0.748 |
| Xiphorhynchus obsoletus | I(time^2):precision0.9 | 4.327 | 0.557 | 3.235 | 5.419 | 7.767 | <.001 |
| Xiphorhynchus obsoletus | I(time^2):precision0.9:recall0.1 | 1.831 | 2.04 | -2.167 | 5.829 | 0.898 | 0.369 |
| Xiphorhynchus obsoletus | I(time^2):precision0.9:recall0.25 | -0.78 | 1.22 | -3.17 | 1.61 | -0.64 | 0.522 |
| Xiphorhynchus obsoletus | I(time^2):precision0.9:recall0.5 | -0.325 | 0.938 | -2.163 | 1.513 | -0.347 | 0.729 |
| Xiphorhynchus obsoletus | I(time^2):precision0.9:recall0.75 | -0.412 | 0.842 | -2.062 | 1.238 | -0.489 | 0.625 |
| Xiphorhynchus obsoletus | I(time^2):precision0.9:recall0.9 | -0.31 | 0.806 | -1.89 | 1.269 | -0.385 | 0.7 |
| Xiphorhynchus obsoletus | I(time^2):recall0.1 | -3.144 | 1.641 | -6.36 | 0.072 | -1.916 | 0.055 |
| Xiphorhynchus obsoletus | I(time^2):recall0.25 | 0.897 | 0.949 | -0.962 | 2.756 | 0.945 | 0.344 |
| Xiphorhynchus obsoletus | I(time^2):recall0.5 | 1.103 | 0.735 | -0.337 | 2.544 | 1.501 | 0.133 |
| Xiphorhynchus obsoletus | I(time^2):recall0.75 | 0.616 | 0.659 | -0.676 | 1.908 | 0.935 | 0.35 |
| Xiphorhynchus obsoletus | I(time^2):recall0.9 | 0.305 | 0.632 | -0.934 | 1.543 | 0.482 | 0.63 |
| Xiphorhynchus obsoletus | precision0.1 | 5.074 | 0.089 | 4.9 | 5.248 | 57.196 | <.001 |
| Xiphorhynchus obsoletus | precision0.1:recall0.1 | 0.13 | 0.315 | -0.488 | 0.747 | 0.412 | 0.68 |
| Xiphorhynchus obsoletus | precision0.1:recall0.25 | -0.201 | 0.196 | -0.586 | 0.183 | -1.027 | 0.304 |
| Xiphorhynchus obsoletus | precision0.1:recall0.5 | -0.197 | 0.151 | -0.493 | 0.099 | -1.307 | 0.191 |
| Xiphorhynchus obsoletus | precision0.1:recall0.75 | -0.105 | 0.134 | -0.369 | 0.158 | -0.783 | 0.434 |
| Xiphorhynchus obsoletus | precision0.1:recall0.9 | -0.051 | 0.128 | -0.303 | 0.201 | -0.398 | 0.691 |
| Xiphorhynchus obsoletus | precision0.25 | 3.819 | 0.091 | 3.641 | 3.997 | 42.049 | <.001 |
| Xiphorhynchus obsoletus | precision0.25:recall0.1 | 0.216 | 0.322 | -0.416 | 0.847 | 0.669 | 0.503 |
| Xiphorhynchus obsoletus | precision0.25:recall0.25 | -0.152 | 0.201 | -0.546 | 0.242 | -0.756 | 0.45 |
| Xiphorhynchus obsoletus | precision0.25:recall0.5 | -0.182 | 0.155 | -0.485 | 0.121 | -1.18 | 0.238 |
| Xiphorhynchus obsoletus | precision0.25:recall0.75 | -0.092 | 0.138 | -0.362 | 0.178 | -0.668 | 0.504 |
| Xiphorhynchus obsoletus | precision0.25:recall0.9 | -0.052 | 0.132 | -0.309 | 0.206 | -0.393 | 0.694 |
| Xiphorhynchus obsoletus | precision0.5 | 2.604 | 0.096 | 2.416 | 2.791 | 27.187 | <.001 |
| Xiphorhynchus obsoletus | precision0.5:recall0.1 | 0.381 | 0.337 | -0.279 | 1.041 | 1.132 | 0.258 |
| Xiphorhynchus obsoletus | precision0.5:recall0.25 | -0.119 | 0.213 | -0.536 | 0.297 | -0.561 | 0.575 |
| Xiphorhynchus obsoletus | precision0.5:recall0.5 | -0.126 | 0.163 | -0.446 | 0.194 | -0.774 | 0.439 |
| Xiphorhynchus obsoletus | precision0.5:recall0.75 | -0.074 | 0.145 | -0.359 | 0.21 | -0.511 | 0.609 |
| Xiphorhynchus obsoletus | precision0.5:recall0.9 | -0.027 | 0.139 | -0.299 | 0.245 | -0.193 | 0.847 |
| Xiphorhynchus obsoletus | precision0.75 | 1.497 | 0.104 | 1.292 | 1.701 | 14.364 | <.001 |
| Xiphorhynchus obsoletus | precision0.75:recall0.1 | 0.28 | 0.365 | -0.436 | 0.996 | 0.767 | 0.443 |
| Xiphorhynchus obsoletus | precision0.75:recall0.25 | -0.029 | 0.231 | -0.482 | 0.423 | -0.127 | 0.899 |
| Xiphorhynchus obsoletus | precision0.75:recall0.5 | -0.004 | 0.177 | -0.351 | 0.343 | -0.022 | 0.982 |
| Xiphorhynchus obsoletus | precision0.75:recall0.75 | -0.037 | 0.158 | -0.347 | 0.273 | -0.233 | 0.816 |
| Xiphorhynchus obsoletus | precision0.75:recall0.9 | -0.031 | 0.151 | -0.327 | 0.265 | -0.207 | 0.836 |
| Xiphorhynchus obsoletus | precision0.9 | 0.749 | 0.113 | 0.528 | 0.97 | 6.646 | <.001 |
| Xiphorhynchus obsoletus | precision0.9:recall0.1 | 0.246 | 0.394 | -0.526 | 1.018 | 0.624 | 0.532 |
| Xiphorhynchus obsoletus | precision0.9:recall0.25 | -0.076 | 0.251 | -0.568 | 0.416 | -0.302 | 0.763 |
| Xiphorhynchus obsoletus | precision0.9:recall0.5 | -0.073 | 0.192 | -0.45 | 0.304 | -0.381 | 0.703 |
| Xiphorhynchus obsoletus | precision0.9:recall0.75 | -0.06 | 0.171 | -0.395 | 0.276 | -0.348 | 0.728 |
| Xiphorhynchus obsoletus | precision0.9:recall0.9 | -0.071 | 0.164 | -0.392 | 0.25 | -0.432 | 0.666 |
| Xiphorhynchus obsoletus | recall0.1 | -2.663 | 0.311 | -3.272 | -2.053 | -8.561 | <.001 |
| Xiphorhynchus obsoletus | recall0.25 | -1.354 | 0.193 | -1.732 | -0.975 | -7.012 | <.001 |
| Xiphorhynchus obsoletus | recall0.5 | -0.584 | 0.148 | -0.875 | -0.294 | -3.94 | <.001 |
| Xiphorhynchus obsoletus | recall0.75 | -0.227 | 0.132 | -0.486 | 0.032 | -1.721 | 0.085 |
| Xiphorhynchus obsoletus | recall0.9 | -0.077 | 0.126 | -0.324 | 0.171 | -0.608 | 0.543 |
| Xiphorhynchus obsoletus | time | 16.487 | 0.401 | 15.701 | 17.272 | 41.145 | <.001 |
| Xiphorhynchus obsoletus | time:precision0.1 | -15.574 | 0.408 | -16.374 | -14.775 | -38.193 | <.001 |
| Xiphorhynchus obsoletus | time:precision0.1:recall0.1 | -2.21 | 1.479 | -5.109 | 0.689 | -1.494 | 0.135 |
| Xiphorhynchus obsoletus | time:precision0.1:recall0.25 | 0.587 | 0.892 | -1.16 | 2.335 | 0.659 | 0.51 |
| Xiphorhynchus obsoletus | time:precision0.1:recall0.5 | 0.697 | 0.689 | -0.653 | 2.047 | 1.012 | 0.312 |
| Xiphorhynchus obsoletus | time:precision0.1:recall0.75 | 0.389 | 0.616 | -0.818 | 1.596 | 0.631 | 0.528 |
| Xiphorhynchus obsoletus | time:precision0.1:recall0.9 | 0.218 | 0.589 | -0.937 | 1.373 | 0.37 | 0.711 |
| Xiphorhynchus obsoletus | time:precision0.25 | -14.129 | 0.417 | -14.947 | -13.312 | -33.875 | <.001 |
| Xiphorhynchus obsoletus | time:precision0.25:recall0.1 | -1.948 | 1.509 | -4.906 | 1.009 | -1.291 | 0.197 |
| Xiphorhynchus obsoletus | time:precision0.25:recall0.25 | 0.753 | 0.914 | -1.038 | 2.544 | 0.824 | 0.41 |
| Xiphorhynchus obsoletus | time:precision0.25:recall0.5 | 0.856 | 0.706 | -0.527 | 2.239 | 1.213 | 0.225 |
| Xiphorhynchus obsoletus | time:precision0.25:recall0.75 | 0.482 | 0.63 | -0.754 | 1.717 | 0.764 | 0.445 |
| Xiphorhynchus obsoletus | time:precision0.25:recall0.9 | 0.253 | 0.603 | -0.929 | 1.435 | 0.419 | 0.675 |
| Xiphorhynchus obsoletus | time:precision0.5 | -11.151 | 0.438 | -12.009 | -10.292 | -25.451 | <.001 |
| Xiphorhynchus obsoletus | time:precision0.5:recall0.1 | -2.492 | 1.571 | -5.572 | 0.588 | -1.586 | 0.113 |
| Xiphorhynchus obsoletus | time:precision0.5:recall0.25 | 0.821 | 0.963 | -1.066 | 2.708 | 0.853 | 0.394 |
| Xiphorhynchus obsoletus | time:precision0.5:recall0.5 | 0.765 | 0.743 | -0.691 | 2.22 | 1.029 | 0.303 |
| Xiphorhynchus obsoletus | time:precision0.5:recall0.75 | 0.356 | 0.663 | -0.943 | 1.655 | 0.538 | 0.591 |
| Xiphorhynchus obsoletus | time:precision0.5:recall0.9 | 0.15 | 0.634 | -1.092 | 1.392 | 0.237 | 0.813 |
| Xiphorhynchus obsoletus | time:precision0.75 | -7.064 | 0.475 | -7.994 | -6.134 | -14.884 | <.001 |
| Xiphorhynchus obsoletus | time:precision0.75:recall0.1 | -1.812 | 1.696 | -5.136 | 1.512 | -1.068 | 0.285 |
| Xiphorhynchus obsoletus | time:precision0.75:recall0.25 | 0.59 | 1.045 | -1.459 | 2.638 | 0.564 | 0.573 |
| Xiphorhynchus obsoletus | time:precision0.75:recall0.5 | 0.228 | 0.804 | -1.346 | 1.803 | 0.284 | 0.776 |
| Xiphorhynchus obsoletus | time:precision0.75:recall0.75 | 0.285 | 0.718 | -1.122 | 1.693 | 0.397 | 0.691 |
| Xiphorhynchus obsoletus | time:precision0.75:recall0.9 | 0.183 | 0.687 | -1.163 | 1.53 | 0.267 | 0.79 |
| Xiphorhynchus obsoletus | time:precision0.9 | -3.65 | 0.515 | -4.66 | -2.64 | -7.086 | <.001 |
| Xiphorhynchus obsoletus | time:precision0.9:recall0.1 | -1.425 | 1.837 | -5.026 | 2.175 | -0.776 | 0.438 |
| Xiphorhynchus obsoletus | time:precision0.9:recall0.25 | 0.561 | 1.137 | -1.667 | 2.789 | 0.493 | 0.622 |
| Xiphorhynchus obsoletus | time:precision0.9:recall0.5 | 0.315 | 0.873 | -1.397 | 2.027 | 0.36 | 0.719 |
| Xiphorhynchus obsoletus | time:precision0.9:recall0.75 | 0.338 | 0.781 | -1.191 | 1.868 | 0.434 | 0.665 |
| Xiphorhynchus obsoletus | time:precision0.9:recall0.9 | 0.314 | 0.747 | -1.15 | 1.777 | 0.42 | 0.674 |
| Xiphorhynchus obsoletus | time:recall0.1 | 2.289 | 1.461 | -0.574 | 5.152 | 1.567 | 0.117 |
| Xiphorhynchus obsoletus | time:recall0.25 | -0.579 | 0.878 | -2.299 | 1.141 | -0.66 | 0.51 |
| Xiphorhynchus obsoletus | time:recall0.5 | -0.836 | 0.677 | -2.163 | 0.491 | -1.235 | 0.217 |
| Xiphorhynchus obsoletus | time:recall0.75 | -0.468 | 0.605 | -1.654 | 0.718 | -0.773 | 0.439 |
| Xiphorhynchus obsoletus | time:recall0.9 | -0.225 | 0.579 | -1.36 | 0.91 | -0.388 | 0.698 |
| Trogon melanurus | (Intercept) | -16.293 | 0.32 | -16.919 | -15.666 | -50.981 | <.001 |
| Trogon melanurus | I(time^2) | -26.223 | 0.659 | -27.514 | -24.932 | -39.819 | <.001 |
| Trogon melanurus | I(time^2):precision0.1 | 25.834 | 0.662 | 24.536 | 27.132 | 39.003 | <.001 |
| Trogon melanurus | I(time^2):precision0.1:recall0.1 | -1.636 | 2.108 | -5.768 | 2.497 | -0.776 | 0.438 |
| Trogon melanurus | I(time^2):precision0.1:recall0.25 | -0.694 | 1.445 | -3.525 | 2.138 | -0.48 | 0.631 |
| Trogon melanurus | I(time^2):precision0.1:recall0.5 | -0.717 | 1.128 | -2.927 | 1.494 | -0.635 | 0.525 |
| Trogon melanurus | I(time^2):precision0.1:recall0.75 | -0.127 | 1.008 | -2.103 | 1.848 | -0.126 | 0.899 |
| Trogon melanurus | I(time^2):precision0.1:recall0.9 | -0.284 | 0.959 | -2.163 | 1.595 | -0.296 | 0.767 |
| Trogon melanurus | I(time^2):precision0.25 | 25.12 | 0.667 | 23.812 | 26.427 | 37.654 | <.001 |
| Trogon melanurus | I(time^2):precision0.25:recall0.1 | -2.161 | 2.126 | -6.329 | 2.006 | -1.016 | 0.309 |
| Trogon melanurus | I(time^2):precision0.25:recall0.25 | -0.882 | 1.456 | -3.736 | 1.971 | -0.606 | 0.544 |
| Trogon melanurus | I(time^2):precision0.25:recall0.5 | -0.874 | 1.136 | -3.101 | 1.353 | -0.769 | 0.442 |
| Trogon melanurus | I(time^2):precision0.25:recall0.75 | -0.177 | 1.015 | -2.166 | 1.813 | -0.174 | 0.862 |
| Trogon melanurus | I(time^2):precision0.25:recall0.9 | -0.236 | 0.966 | -2.128 | 1.657 | -0.244 | 0.807 |
| Trogon melanurus | I(time^2):precision0.5 | 22.894 | 0.678 | 21.564 | 24.224 | 33.743 | <.001 |
| Trogon melanurus | I(time^2):precision0.5:recall0.1 | -1.69 | 2.165 | -5.932 | 2.553 | -0.781 | 0.435 |
| Trogon melanurus | I(time^2):precision0.5:recall0.25 | -0.995 | 1.483 | -3.901 | 1.912 | -0.671 | 0.502 |
| Trogon melanurus | I(time^2):precision0.5:recall0.5 | -0.583 | 1.156 | -2.848 | 1.683 | -0.504 | 0.614 |
| Trogon melanurus | I(time^2):precision0.5:recall0.75 | -0.088 | 1.033 | -2.112 | 1.936 | -0.085 | 0.932 |
| Trogon melanurus | I(time^2):precision0.5:recall0.9 | -0.193 | 0.982 | -2.118 | 1.732 | -0.196 | 0.845 |
| Trogon melanurus | I(time^2):precision0.75 | 18.982 | 0.705 | 17.601 | 20.363 | 26.943 | <.001 |
| Trogon melanurus | I(time^2):precision0.75:recall0.1 | -1.29 | 2.248 | -5.697 | 3.117 | -0.574 | 0.566 |
| Trogon melanurus | I(time^2):precision0.75:recall0.25 | -2.272 | 1.556 | -5.322 | 0.778 | -1.46 | 0.144 |
| Trogon melanurus | I(time^2):precision0.75:recall0.5 | -0.546 | 1.201 | -2.9 | 1.808 | -0.455 | 0.649 |
| Trogon melanurus | I(time^2):precision0.75:recall0.75 | 0.018 | 1.072 | -2.083 | 2.118 | 0.016 | 0.987 |
| Trogon melanurus | I(time^2):precision0.75:recall0.9 | -0.213 | 1.02 | -2.213 | 1.786 | -0.209 | 0.834 |
| Trogon melanurus | I(time^2):precision0.9 | 12.095 | 0.771 | 10.584 | 13.606 | 15.687 | <.001 |
| Trogon melanurus | I(time^2):precision0.9:recall0.1 | -1.908 | 2.484 | -6.777 | 2.961 | -0.768 | 0.442 |
| Trogon melanurus | I(time^2):precision0.9:recall0.25 | -2.89 | 1.728 | -6.277 | 0.497 | -1.672 | 0.094 |
| Trogon melanurus | I(time^2):precision0.9:recall0.5 | 0.741 | 1.302 | -1.81 | 3.293 | 0.57 | 0.569 |
| Trogon melanurus | I(time^2):precision0.9:recall0.75 | 0.459 | 1.169 | -1.832 | 2.751 | 0.393 | 0.694 |
| Trogon melanurus | I(time^2):precision0.9:recall0.9 | 0.337 | 1.113 | -1.844 | 2.518 | 0.303 | 0.762 |
| Trogon melanurus | I(time^2):recall0.1 | 1.554 | 2.097 | -2.556 | 5.665 | 0.741 | 0.459 |
| Trogon melanurus | I(time^2):recall0.25 | 0.753 | 1.437 | -2.063 | 3.57 | 0.524 | 0.6 |
| Trogon melanurus | I(time^2):recall0.5 | 0.79 | 1.122 | -1.408 | 2.989 | 0.705 | 0.481 |
| Trogon melanurus | I(time^2):recall0.75 | 0.188 | 1.002 | -1.776 | 2.153 | 0.188 | 0.851 |
| Trogon melanurus | I(time^2):recall0.9 | 0.224 | 0.953 | -1.644 | 2.092 | 0.235 | 0.814 |
| Trogon melanurus | precision0.1 | 14.78 | 0.32 | 14.153 | 15.407 | 46.187 | <.001 |
| Trogon melanurus | precision0.1:recall0.1 | -1.011 | 1.011 | -2.993 | 0.971 | -0.999 | 0.318 |
| Trogon melanurus | precision0.1:recall0.25 | -0.646 | 0.693 | -2.004 | 0.711 | -0.933 | 0.351 |
| Trogon melanurus | precision0.1:recall0.5 | -0.524 | 0.543 | -1.588 | 0.54 | -0.966 | 0.334 |
| Trogon melanurus | precision0.1:recall0.75 | -0.164 | 0.486 | -1.117 | 0.789 | -0.338 | 0.736 |
| Trogon melanurus | precision0.1:recall0.9 | -0.159 | 0.463 | -1.066 | 0.748 | -0.343 | 0.732 |
| Trogon melanurus | precision0.25 | 13.376 | 0.321 | 12.747 | 14.004 | 41.709 | <.001 |
| Trogon melanurus | precision0.25:recall0.1 | -1.049 | 1.014 | -3.037 | 0.938 | -1.035 | 0.301 |
| Trogon melanurus | precision0.25:recall0.25 | -0.587 | 0.694 | -1.947 | 0.774 | -0.845 | 0.398 |
| Trogon melanurus | precision0.25:recall0.5 | -0.484 | 0.544 | -1.55 | 0.582 | -0.889 | 0.374 |
| Trogon melanurus | precision0.25:recall0.75 | -0.132 | 0.487 | -1.087 | 0.823 | -0.271 | 0.786 |
| Trogon melanurus | precision0.25:recall0.9 | -0.131 | 0.464 | -1.04 | 0.778 | -0.283 | 0.777 |
| Trogon melanurus | precision0.5 | 11.703 | 0.323 | 11.069 | 12.336 | 36.21 | <.001 |
| Trogon melanurus | precision0.5:recall0.1 | -0.788 | 1.022 | -2.792 | 1.216 | -0.771 | 0.441 |
| Trogon melanurus | precision0.5:recall0.25 | -0.664 | 0.701 | -2.037 | 0.709 | -0.948 | 0.343 |
| Trogon melanurus | precision0.5:recall0.5 | -0.386 | 0.548 | -1.46 | 0.689 | -0.703 | 0.482 |
| Trogon melanurus | precision0.5:recall0.75 | -0.1 | 0.491 | -1.063 | 0.863 | -0.204 | 0.839 |
| Trogon melanurus | precision0.5:recall0.9 | -0.101 | 0.467 | -1.017 | 0.815 | -0.217 | 0.828 |
| Trogon melanurus | precision0.75 | 9.538 | 0.332 | 8.887 | 10.189 | 28.719 | <.001 |
| Trogon melanurus | precision0.75:recall0.1 | -0.618 | 1.05 | -2.677 | 1.44 | -0.589 | 0.556 |
| Trogon melanurus | precision0.75:recall0.25 | -1.166 | 0.727 | -2.592 | 0.26 | -1.603 | 0.109 |
| Trogon melanurus | precision0.75:recall0.5 | -0.383 | 0.564 | -1.488 | 0.722 | -0.679 | 0.497 |
| Trogon melanurus | precision0.75:recall0.75 | -0.051 | 0.505 | -1.04 | 0.938 | -0.101 | 0.92 |
| Trogon melanurus | precision0.75:recall0.9 | -0.133 | 0.481 | -1.075 | 0.809 | -0.277 | 0.782 |
| Trogon melanurus | precision0.9 | 6.137 | 0.363 | 5.425 | 6.849 | 16.892 | <.001 |
| Trogon melanurus | precision0.9:recall0.1 | -0.813 | 1.16 | -3.087 | 1.46 | -0.701 | 0.483 |
| Trogon melanurus | precision0.9:recall0.25 | -1.452 | 0.811 | -3.041 | 0.137 | -1.791 | 0.073 |
| Trogon melanurus | precision0.9:recall0.5 | 0.261 | 0.61 | -0.936 | 1.457 | 0.427 | 0.669 |
| Trogon melanurus | precision0.9:recall0.75 | 0.163 | 0.55 | -0.915 | 1.241 | 0.296 | 0.767 |
| Trogon melanurus | precision0.9:recall0.9 | 0.133 | 0.524 | -0.894 | 1.16 | 0.254 | 0.8 |
| Trogon melanurus | recall0.1 | -1.514 | 1.01 | -3.493 | 0.466 | -1.499 | 0.134 |
| Trogon melanurus | recall0.25 | -0.886 | 0.692 | -2.242 | 0.47 | -1.281 | 0.2 |
| Trogon melanurus | recall0.5 | -0.253 | 0.542 | -1.315 | 0.81 | -0.466 | 0.641 |
| Trogon melanurus | recall0.75 | -0.165 | 0.486 | -1.117 | 0.787 | -0.34 | 0.734 |
| Trogon melanurus | recall0.9 | 0.018 | 0.462 | -0.887 | 0.924 | 0.04 | 0.968 |
| Trogon melanurus | time | 37.736 | 0.926 | 35.921 | 39.551 | 40.748 | <.001 |
| Trogon melanurus | time:precision0.1 | -37.059 | 0.929 | -38.88 | -35.239 | -39.89 | <.001 |
| Trogon melanurus | time:precision0.1:recall0.1 | 2.367 | 2.947 | -3.408 | 8.142 | 0.803 | 0.422 |
| Trogon melanurus | time:precision0.1:recall0.25 | 1.191 | 2.018 | -2.765 | 5.146 | 0.59 | 0.555 |
| Trogon melanurus | time:precision0.1:recall0.5 | 1.098 | 1.579 | -1.997 | 4.192 | 0.695 | 0.487 |
| Trogon melanurus | time:precision0.1:recall0.75 | 0.237 | 1.413 | -2.532 | 3.006 | 0.168 | 0.867 |
| Trogon melanurus | time:precision0.1:recall0.9 | 0.402 | 1.344 | -2.233 | 3.036 | 0.299 | 0.765 |
| Trogon melanurus | time:precision0.25 | -35.943 | 0.933 | -37.772 | -34.114 | -38.519 | <.001 |
| Trogon melanurus | time:precision0.25:recall0.1 | 3.046 | 2.962 | -2.76 | 8.852 | 1.028 | 0.304 |
| Trogon melanurus | time:precision0.25:recall0.25 | 1.427 | 2.028 | -2.548 | 5.402 | 0.704 | 0.482 |
| Trogon melanurus | time:precision0.25:recall0.5 | 1.284 | 1.586 | -1.825 | 4.393 | 0.81 | 0.418 |
| Trogon melanurus | time:precision0.25:recall0.75 | 0.292 | 1.419 | -2.489 | 3.073 | 0.206 | 0.837 |
| Trogon melanurus | time:precision0.25:recall0.9 | 0.35 | 1.35 | -2.296 | 2.996 | 0.259 | 0.795 |
| Trogon melanurus | time:precision0.5 | -32.773 | 0.945 | -34.624 | -30.922 | -34.699 | <.001 |
| Trogon melanurus | time:precision0.5:recall0.1 | 2.328 | 3 | -3.552 | 8.208 | 0.776 | 0.438 |
| Trogon melanurus | time:precision0.5:recall0.25 | 1.685 | 2.056 | -2.344 | 5.714 | 0.82 | 0.412 |
| Trogon melanurus | time:precision0.5:recall0.5 | 0.953 | 1.606 | -2.194 | 4.1 | 0.594 | 0.553 |
| Trogon melanurus | time:precision0.5:recall0.75 | 0.197 | 1.436 | -2.618 | 3.013 | 0.137 | 0.891 |
| Trogon melanurus | time:precision0.5:recall0.9 | 0.277 | 1.367 | -2.401 | 2.956 | 0.203 | 0.839 |
| Trogon melanurus | time:precision0.75 | -27.219 | 0.976 | -29.133 | -25.306 | -27.883 | <.001 |
| Trogon melanurus | time:precision0.75:recall0.1 | 1.796 | 3.101 | -4.282 | 7.873 | 0.579 | 0.563 |
| Trogon melanurus | time:precision0.75:recall0.25 | 3.369 | 2.148 | -0.84 | 7.578 | 1.569 | 0.117 |
| Trogon melanurus | time:precision0.75:recall0.5 | 0.939 | 1.661 | -2.316 | 4.194 | 0.566 | 0.572 |
| Trogon melanurus | time:precision0.75:recall0.75 | 0.052 | 1.484 | -2.856 | 2.961 | 0.035 | 0.972 |
| Trogon melanurus | time:precision0.75:recall0.9 | 0.347 | 1.413 | -2.423 | 3.116 | 0.245 | 0.806 |
| Trogon melanurus | time:precision0.9 | -17.458 | 1.069 | -19.553 | -15.362 | -16.326 | <.001 |
| Trogon melanurus | time:precision0.9:recall0.1 | 2.549 | 3.429 | -4.172 | 9.27 | 0.743 | 0.457 |
| Trogon melanurus | time:precision0.9:recall0.25 | 4.2 | 2.391 | -0.487 | 8.887 | 1.756 | 0.079 |
| Trogon melanurus | time:precision0.9:recall0.5 | -0.931 | 1.801 | -4.46 | 2.599 | -0.517 | 0.605 |
| Trogon melanurus | time:precision0.9:recall0.75 | -0.576 | 1.62 | -3.752 | 2.6 | -0.355 | 0.722 |
| Trogon melanurus | time:precision0.9:recall0.9 | -0.442 | 1.543 | -3.466 | 2.581 | -0.287 | 0.774 |
| Trogon melanurus | time:recall0.1 | -2.318 | 2.938 | -8.076 | 3.441 | -0.789 | 0.43 |
| Trogon melanurus | time:recall0.25 | -1.301 | 2.012 | -5.245 | 2.643 | -0.646 | 0.518 |
| Trogon melanurus | time:recall0.5 | -1.234 | 1.574 | -4.318 | 1.851 | -0.784 | 0.433 |
| Trogon melanurus | time:recall0.75 | -0.33 | 1.408 | -3.09 | 2.43 | -0.234 | 0.815 |
| Trogon melanurus | time:recall0.9 | -0.345 | 1.34 | -2.971 | 2.281 | -0.257 | 0.797 |
| Turdus albicollis | (Intercept) | -3.762 | 0.054 | -3.868 | -3.657 | -70.048 | <.001 |
| Turdus albicollis | I(time^2) | -41.64 | 1.351 | -44.288 | -38.993 | -30.822 | <.001 |
| Turdus albicollis | I(time^2):precision0.1 | 42.064 | 1.353 | 39.412 | 44.716 | 31.088 | <.001 |
| Turdus albicollis | I(time^2):precision0.1:recall0.1 | -2.553 | 4.314 | -11.008 | 5.902 | -0.592 | 0.554 |
| Turdus albicollis | I(time^2):precision0.1:recall0.25 | -0.91 | 2.991 | -6.773 | 4.953 | -0.304 | 0.761 |
| Turdus albicollis | I(time^2):precision0.1:recall0.5 | 0.533 | 2.347 | -4.067 | 5.134 | 0.227 | 0.82 |
| Turdus albicollis | I(time^2):precision0.1:recall0.75 | -0.474 | 2.064 | -4.519 | 3.572 | -0.229 | 0.818 |
| Turdus albicollis | I(time^2):precision0.1:recall0.9 | 0.201 | 1.974 | -3.667 | 4.07 | 0.102 | 0.919 |
| Turdus albicollis | I(time^2):precision0.25 | 42.502 | 1.356 | 39.845 | 45.159 | 31.352 | <.001 |
| Turdus albicollis | I(time^2):precision0.25:recall0.1 | -2.619 | 4.323 | -11.092 | 5.854 | -0.606 | 0.545 |
| Turdus albicollis | I(time^2):precision0.25:recall0.25 | -1.089 | 2.997 | -6.964 | 4.785 | -0.363 | 0.716 |
| Turdus albicollis | I(time^2):precision0.25:recall0.5 | 0.759 | 2.352 | -3.851 | 5.368 | 0.323 | 0.747 |
| Turdus albicollis | I(time^2):precision0.25:recall0.75 | -0.335 | 2.068 | -4.388 | 3.718 | -0.162 | 0.871 |
| Turdus albicollis | I(time^2):precision0.25:recall0.9 | 0.264 | 1.978 | -3.612 | 4.14 | 0.133 | 0.894 |
| Turdus albicollis | I(time^2):precision0.5 | 42.672 | 1.361 | 40.004 | 45.339 | 31.352 | <.001 |
| Turdus albicollis | I(time^2):precision0.5:recall0.1 | -2.731 | 4.342 | -11.242 | 5.78 | -0.629 | 0.529 |
| Turdus albicollis | I(time^2):precision0.5:recall0.25 | -1.511 | 3.01 | -7.411 | 4.388 | -0.502 | 0.616 |
| Turdus albicollis | I(time^2):precision0.5:recall0.5 | 0.175 | 2.361 | -4.453 | 4.803 | 0.074 | 0.941 |
| Turdus albicollis | I(time^2):precision0.5:recall0.75 | -0.425 | 2.076 | -4.494 | 3.645 | -0.204 | 0.838 |
| Turdus albicollis | I(time^2):precision0.5:recall0.9 | 0.228 | 1.985 | -3.663 | 4.119 | 0.115 | 0.909 |
| Turdus albicollis | I(time^2):precision0.75 | 40.816 | 1.373 | 38.125 | 43.508 | 29.719 | <.001 |
| Turdus albicollis | I(time^2):precision0.75:recall0.1 | -3.444 | 4.388 | -12.045 | 5.157 | -0.785 | 0.433 |
| Turdus albicollis | I(time^2):precision0.75:recall0.25 | -1.132 | 3.037 | -7.084 | 4.821 | -0.373 | 0.709 |
| Turdus albicollis | I(time^2):precision0.75:recall0.5 | -0.143 | 2.384 | -4.816 | 4.53 | -0.06 | 0.952 |
| Turdus albicollis | I(time^2):precision0.75:recall0.75 | -0.625 | 2.096 | -4.732 | 3.483 | -0.298 | 0.766 |
| Turdus albicollis | I(time^2):precision0.75:recall0.9 | 0.359 | 2.003 | -3.567 | 4.286 | 0.179 | 0.858 |
| Turdus albicollis | I(time^2):precision0.9 | 35.005 | 1.42 | 32.222 | 37.789 | 24.651 | <.001 |
| Turdus albicollis | I(time^2):precision0.9:recall0.1 | -7.824 | 4.692 | -17.021 | 1.373 | -1.667 | 0.095 |
| Turdus albicollis | I(time^2):precision0.9:recall0.25 | 0.268 | 3.122 | -5.851 | 6.386 | 0.086 | 0.932 |
| Turdus albicollis | I(time^2):precision0.9:recall0.5 | -1.317 | 2.481 | -6.18 | 3.546 | -0.531 | 0.596 |
| Turdus albicollis | I(time^2):precision0.9:recall0.75 | -0.47 | 2.168 | -4.719 | 3.779 | -0.217 | 0.828 |
| Turdus albicollis | I(time^2):precision0.9:recall0.9 | 0.356 | 2.071 | -3.703 | 4.414 | 0.172 | 0.864 |
| Turdus albicollis | I(time^2):recall0.1 | 2.355 | 4.308 | -6.088 | 10.799 | 0.547 | 0.585 |
| Turdus albicollis | I(time^2):recall0.25 | 0.893 | 2.987 | -4.962 | 6.748 | 0.299 | 0.765 |
| Turdus albicollis | I(time^2):recall0.5 | -0.712 | 2.344 | -5.306 | 3.881 | -0.304 | 0.761 |
| Turdus albicollis | I(time^2):recall0.75 | 0.406 | 2.061 | -3.633 | 4.445 | 0.197 | 0.844 |
| Turdus albicollis | I(time^2):recall0.9 | -0.223 | 1.971 | -4.086 | 3.639 | -0.113 | 0.91 |
| Turdus albicollis | precision0.1 | 2.479 | 0.056 | 2.369 | 2.588 | 44.194 | <.001 |
| Turdus albicollis | precision0.1:recall0.1 | -0.182 | 0.183 | -0.54 | 0.177 | -0.994 | 0.32 |
| Turdus albicollis | precision0.1:recall0.25 | -0.172 | 0.124 | -0.414 | 0.07 | -1.393 | 0.164 |
| Turdus albicollis | precision0.1:recall0.5 | -0.066 | 0.098 | -0.257 | 0.126 | -0.673 | 0.501 |
| Turdus albicollis | precision0.1:recall0.75 | -0.097 | 0.085 | -0.264 | 0.07 | -1.139 | 0.255 |
| Turdus albicollis | precision0.1:recall0.9 | -0.041 | 0.081 | -0.2 | 0.119 | -0.499 | 0.618 |
| Turdus albicollis | precision0.25 | 1.762 | 0.058 | 1.649 | 1.876 | 30.413 | <.001 |
| Turdus albicollis | precision0.25:recall0.1 | -0.124 | 0.19 | -0.496 | 0.248 | -0.655 | 0.512 |
| Turdus albicollis | precision0.25:recall0.25 | -0.064 | 0.128 | -0.314 | 0.186 | -0.501 | 0.616 |
| Turdus albicollis | precision0.25:recall0.5 | 0.057 | 0.101 | -0.141 | 0.254 | 0.565 | 0.572 |
| Turdus albicollis | precision0.25:recall0.75 | -0.041 | 0.088 | -0.213 | 0.131 | -0.465 | 0.642 |
| Turdus albicollis | precision0.25:recall0.9 | -0.016 | 0.084 | -0.181 | 0.148 | -0.196 | 0.845 |
| Turdus albicollis | precision0.5 | 1.457 | 0.06 | 1.34 | 1.574 | 24.312 | <.001 |
| Turdus albicollis | precision0.5:recall0.1 | -0.067 | 0.196 | -0.452 | 0.317 | -0.343 | 0.732 |
| Turdus albicollis | precision0.5:recall0.25 | -0.106 | 0.133 | -0.366 | 0.154 | -0.8 | 0.424 |
| Turdus albicollis | precision0.5:recall0.5 | 0.008 | 0.104 | -0.197 | 0.212 | 0.072 | 0.943 |
| Turdus albicollis | precision0.5:recall0.75 | -0.031 | 0.091 | -0.209 | 0.147 | -0.342 | 0.733 |
| Turdus albicollis | precision0.5:recall0.9 | -0.015 | 0.087 | -0.186 | 0.155 | -0.174 | 0.862 |
| Turdus albicollis | precision0.75 | 1.269 | 0.062 | 1.148 | 1.39 | 20.557 | <.001 |
| Turdus albicollis | precision0.75:recall0.1 | -0.142 | 0.203 | -0.541 | 0.257 | -0.698 | 0.485 |
| Turdus albicollis | precision0.75:recall0.25 | -0.072 | 0.137 | -0.34 | 0.195 | -0.528 | 0.597 |
| Turdus albicollis | precision0.75:recall0.5 | -0.012 | 0.108 | -0.223 | 0.199 | -0.115 | 0.908 |
| Turdus albicollis | precision0.75:recall0.75 | -0.036 | 0.094 | -0.219 | 0.148 | -0.379 | 0.705 |
| Turdus albicollis | precision0.75:recall0.9 | 0 | 0.09 | -0.176 | 0.175 | -0.003 | 0.997 |
| Turdus albicollis | precision0.9 | 1.014 | 0.064 | 0.888 | 1.14 | 15.776 | <.001 |
| Turdus albicollis | precision0.9:recall0.1 | -0.27 | 0.217 | -0.696 | 0.155 | -1.246 | 0.213 |
| Turdus albicollis | precision0.9:recall0.25 | -0.003 | 0.141 | -0.28 | 0.274 | -0.022 | 0.982 |
| Turdus albicollis | precision0.9:recall0.5 | -0.04 | 0.113 | -0.261 | 0.181 | -0.355 | 0.723 |
| Turdus albicollis | precision0.9:recall0.75 | -0.019 | 0.098 | -0.21 | 0.173 | -0.192 | 0.847 |
| Turdus albicollis | precision0.9:recall0.9 | 0.004 | 0.093 | -0.179 | 0.186 | 0.038 | 0.969 |
| Turdus albicollis | recall0.1 | -2.349 | 0.176 | -2.695 | -2.004 | -13.331 | <.001 |
| Turdus albicollis | recall0.25 | -1.406 | 0.119 | -1.639 | -1.173 | -11.832 | <.001 |
| Turdus albicollis | recall0.5 | -0.783 | 0.094 | -0.967 | -0.6 | -8.355 | <.001 |
| Turdus albicollis | recall0.75 | -0.268 | 0.082 | -0.428 | -0.108 | -3.288 | 0.001 |
| Turdus albicollis | recall0.9 | -0.102 | 0.078 | -0.255 | 0.051 | -1.305 | 0.192 |
| Turdus albicollis | time | 13.827 | 0.568 | 12.713 | 14.94 | 24.337 | <.001 |
| Turdus albicollis | time:precision0.1 | -14.605 | 0.573 | -15.729 | -13.481 | -25.477 | <.001 |
| Turdus albicollis | time:precision0.1:recall0.1 | 0.833 | 1.851 | -2.794 | 4.46 | 0.45 | 0.653 |
| Turdus albicollis | time:precision0.1:recall0.25 | 0.423 | 1.266 | -2.058 | 2.904 | 0.334 | 0.739 |
| Turdus albicollis | time:precision0.1:recall0.5 | -0.281 | 0.995 | -2.233 | 1.67 | -0.283 | 0.777 |
| Turdus albicollis | time:precision0.1:recall0.75 | 0.368 | 0.872 | -1.341 | 2.078 | 0.422 | 0.673 |
| Turdus albicollis | time:precision0.1:recall0.9 | 0.048 | 0.834 | -1.586 | 1.683 | 0.058 | 0.954 |
| Turdus albicollis | time:precision0.25 | -15.599 | 0.579 | -16.733 | -14.464 | -26.951 | <.001 |
| Turdus albicollis | time:precision0.25:recall0.1 | 0.995 | 1.87 | -2.67 | 4.661 | 0.532 | 0.595 |
| Turdus albicollis | time:precision0.25:recall0.25 | 0.505 | 1.278 | -2.001 | 3.011 | 0.395 | 0.693 |
| Turdus albicollis | time:precision0.25:recall0.5 | -0.554 | 1.005 | -2.523 | 1.416 | -0.551 | 0.582 |
| Turdus albicollis | time:precision0.25:recall0.75 | 0.231 | 0.881 | -1.495 | 1.958 | 0.263 | 0.793 |
| Turdus albicollis | time:precision0.25:recall0.9 | -0.013 | 0.842 | -1.663 | 1.638 | -0.015 | 0.988 |
| Turdus albicollis | time:precision0.5 | -16.803 | 0.588 | -17.956 | -15.651 | -28.573 | <.001 |
| Turdus albicollis | time:precision0.5:recall0.1 | 1.004 | 1.902 | -2.724 | 4.732 | 0.528 | 0.598 |
| Turdus albicollis | time:precision0.5:recall0.25 | 0.927 | 1.3 | -1.621 | 3.475 | 0.713 | 0.476 |
| Turdus albicollis | time:precision0.5:recall0.5 | -0.043 | 1.021 | -2.045 | 1.959 | -0.042 | 0.966 |
| Turdus albicollis | time:precision0.5:recall0.75 | 0.282 | 0.895 | -1.472 | 2.036 | 0.315 | 0.752 |
| Turdus albicollis | time:precision0.5:recall0.9 | 0.017 | 0.856 | -1.66 | 1.694 | 0.02 | 0.984 |
| Turdus albicollis | time:precision0.75 | -16.642 | 0.602 | -17.822 | -15.462 | -27.641 | <.001 |
| Turdus albicollis | time:precision0.75:recall0.1 | 1.657 | 1.952 | -2.169 | 5.483 | 0.849 | 0.396 |
| Turdus albicollis | time:precision0.75:recall0.25 | 0.658 | 1.331 | -1.95 | 3.266 | 0.494 | 0.621 |
| Turdus albicollis | time:precision0.75:recall0.5 | 0.177 | 1.047 | -1.875 | 2.228 | 0.169 | 0.866 |
| Turdus albicollis | time:precision0.75:recall0.75 | 0.391 | 0.917 | -1.406 | 2.188 | 0.427 | 0.67 |
| Turdus albicollis | time:precision0.75:recall0.9 | -0.091 | 0.876 | -1.807 | 1.626 | -0.104 | 0.918 |
| Turdus albicollis | time:precision0.9 | -14.113 | 0.632 | -15.351 | -12.874 | -22.34 | <.001 |
| Turdus albicollis | time:precision0.9:recall0.1 | 3.612 | 2.119 | -0.541 | 7.765 | 1.704 | 0.088 |
| Turdus albicollis | time:precision0.9:recall0.25 | -0.108 | 1.387 | -2.826 | 2.61 | -0.078 | 0.938 |
| Turdus albicollis | time:precision0.9:recall0.5 | 0.657 | 1.105 | -1.51 | 2.823 | 0.594 | 0.552 |
| Turdus albicollis | time:precision0.9:recall0.75 | 0.239 | 0.962 | -1.647 | 2.126 | 0.249 | 0.803 |
| Turdus albicollis | time:precision0.9:recall0.9 | -0.108 | 0.919 | -1.909 | 1.693 | -0.117 | 0.907 |
| Turdus albicollis | time:recall0.1 | -0.631 | 1.836 | -4.229 | 2.968 | -0.343 | 0.731 |
| Turdus albicollis | time:recall0.25 | -0.334 | 1.256 | -2.795 | 2.126 | -0.266 | 0.79 |
| Turdus albicollis | time:recall0.5 | 0.501 | 0.987 | -1.433 | 2.436 | 0.508 | 0.611 |
| Turdus albicollis | time:recall0.75 | -0.272 | 0.865 | -1.967 | 1.423 | -0.315 | 0.753 |
| Turdus albicollis | time:recall0.9 | -0.006 | 0.827 | -1.626 | 1.615 | -0.007 | 0.995 |
| Crypturellus undulatus | (Intercept) | -3.993 | 0.041 | -4.074 | -3.913 | -97.486 | <.001 |
| Crypturellus undulatus | I(time^2) | -2.568 | 0.158 | -2.878 | -2.258 | -16.232 | <.001 |
| Crypturellus undulatus | I(time^2):precision0.1 | 2.37 | 0.169 | 2.04 | 2.701 | 14.042 | <.001 |
| Crypturellus undulatus | I(time^2):precision0.1:recall0.1 | -0.145 | 0.541 | -1.206 | 0.916 | -0.268 | 0.789 |
| Crypturellus undulatus | I(time^2):precision0.1:recall0.25 | 0.125 | 0.37 | -0.6 | 0.85 | 0.338 | 0.735 |
| Crypturellus undulatus | I(time^2):precision0.1:recall0.5 | -0.015 | 0.288 | -0.579 | 0.549 | -0.052 | 0.959 |
| Crypturellus undulatus | I(time^2):precision0.1:recall0.75 | 0.103 | 0.256 | -0.399 | 0.605 | 0.401 | 0.688 |
| Crypturellus undulatus | I(time^2):precision0.1:recall0.9 | -0.006 | 0.244 | -0.485 | 0.473 | -0.025 | 0.98 |
| Crypturellus undulatus | I(time^2):precision0.25 | 2.018 | 0.177 | 1.671 | 2.364 | 11.404 | <.001 |
| Crypturellus undulatus | I(time^2):precision0.25:recall0.1 | -0.029 | 0.573 | -1.151 | 1.094 | -0.05 | 0.96 |
| Crypturellus undulatus | I(time^2):precision0.25:recall0.25 | 0.061 | 0.391 | -0.705 | 0.827 | 0.156 | 0.876 |
| Crypturellus undulatus | I(time^2):precision0.25:recall0.5 | -0.131 | 0.303 | -0.725 | 0.464 | -0.431 | 0.666 |
| Crypturellus undulatus | I(time^2):precision0.25:recall0.75 | 0.06 | 0.269 | -0.467 | 0.588 | 0.224 | 0.823 |
| Crypturellus undulatus | I(time^2):precision0.25:recall0.9 | -0.009 | 0.256 | -0.512 | 0.493 | -0.037 | 0.971 |
| Crypturellus undulatus | I(time^2):precision0.5 | 1.309 | 0.192 | 0.933 | 1.686 | 6.813 | <.001 |
| Crypturellus undulatus | I(time^2):precision0.5:recall0.1 | 0.435 | 0.623 | -0.786 | 1.656 | 0.698 | 0.485 |
| Crypturellus undulatus | I(time^2):precision0.5:recall0.25 | 0.224 | 0.425 | -0.61 | 1.058 | 0.527 | 0.598 |
| Crypturellus undulatus | I(time^2):precision0.5:recall0.5 | -0.041 | 0.33 | -0.689 | 0.606 | -0.125 | 0.9 |
| Crypturellus undulatus | I(time^2):precision0.5:recall0.75 | 0.089 | 0.293 | -0.485 | 0.663 | 0.303 | 0.762 |
| Crypturellus undulatus | I(time^2):precision0.5:recall0.9 | 0.024 | 0.279 | -0.522 | 0.57 | 0.087 | 0.931 |
| Crypturellus undulatus | I(time^2):precision0.75 | 0.661 | 0.208 | 0.254 | 1.068 | 3.182 | 0.001 |
| Crypturellus undulatus | I(time^2):precision0.75:recall0.1 | 0.285 | 0.676 | -1.04 | 1.609 | 0.421 | 0.674 |
| Crypturellus undulatus | I(time^2):precision0.75:recall0.25 | 0.157 | 0.461 | -0.746 | 1.06 | 0.34 | 0.734 |
| Crypturellus undulatus | I(time^2):precision0.75:recall0.5 | -0.019 | 0.357 | -0.719 | 0.682 | -0.052 | 0.959 |
| Crypturellus undulatus | I(time^2):precision0.75:recall0.75 | 0.073 | 0.317 | -0.547 | 0.694 | 0.231 | 0.817 |
| Crypturellus undulatus | I(time^2):precision0.75:recall0.9 | 0.05 | 0.301 | -0.541 | 0.64 | 0.166 | 0.869 |
| Crypturellus undulatus | I(time^2):precision0.9 | 0.275 | 0.217 | -0.15 | 0.701 | 1.267 | 0.205 |
| Crypturellus undulatus | I(time^2):precision0.9:recall0.1 | 0.048 | 0.709 | -1.342 | 1.437 | 0.067 | 0.946 |
| Crypturellus undulatus | I(time^2):precision0.9:recall0.25 | 0.184 | 0.482 | -0.762 | 1.13 | 0.381 | 0.703 |
| Crypturellus undulatus | I(time^2):precision0.9:recall0.5 | 0.051 | 0.374 | -0.682 | 0.783 | 0.135 | 0.892 |
| Crypturellus undulatus | I(time^2):precision0.9:recall0.75 | 0.034 | 0.331 | -0.615 | 0.684 | 0.104 | 0.917 |
| Crypturellus undulatus | I(time^2):precision0.9:recall0.9 | 0.064 | 0.315 | -0.553 | 0.681 | 0.203 | 0.839 |
| Crypturellus undulatus | I(time^2):recall0.1 | 0.11 | 0.518 | -0.905 | 1.124 | 0.212 | 0.832 |
| Crypturellus undulatus | I(time^2):recall0.25 | -0.174 | 0.352 | -0.865 | 0.516 | -0.494 | 0.621 |
| Crypturellus undulatus | I(time^2):recall0.5 | 0.054 | 0.272 | -0.48 | 0.588 | 0.197 | 0.844 |
| Crypturellus undulatus | I(time^2):recall0.75 | -0.058 | 0.241 | -0.531 | 0.415 | -0.24 | 0.81 |
| Crypturellus undulatus | I(time^2):recall0.9 | 0.04 | 0.229 | -0.409 | 0.49 | 0.176 | 0.86 |
| Crypturellus undulatus | precision0.1 | 3.547 | 0.043 | 3.463 | 3.632 | 82.411 | <.001 |
| Crypturellus undulatus | precision0.1:recall0.1 | -0.441 | 0.14 | -0.715 | -0.167 | -3.153 | 0.002 |
| Crypturellus undulatus | precision0.1:recall0.25 | -0.335 | 0.096 | -0.522 | -0.148 | -3.51 | <.001 |
| Crypturellus undulatus | precision0.1:recall0.5 | -0.259 | 0.074 | -0.403 | -0.114 | -3.503 | <.001 |
| Crypturellus undulatus | precision0.1:recall0.75 | -0.128 | 0.065 | -0.256 | 0.001 | -1.951 | 0.051 |
| Crypturellus undulatus | precision0.1:recall0.9 | -0.066 | 0.062 | -0.189 | 0.056 | -1.066 | 0.286 |
| Crypturellus undulatus | precision0.25 | 2.187 | 0.045 | 2.099 | 2.275 | 48.818 | <.001 |
| Crypturellus undulatus | precision0.25:recall0.1 | -0.084 | 0.146 | -0.371 | 0.202 | -0.577 | 0.564 |
| Crypturellus undulatus | precision0.25:recall0.25 | -0.088 | 0.1 | -0.284 | 0.108 | -0.881 | 0.378 |
| Crypturellus undulatus | precision0.25:recall0.5 | -0.089 | 0.077 | -0.24 | 0.062 | -1.157 | 0.247 |
| Crypturellus undulatus | precision0.25:recall0.75 | -0.023 | 0.068 | -0.157 | 0.111 | -0.336 | 0.737 |
| Crypturellus undulatus | precision0.25:recall0.9 | -0.018 | 0.065 | -0.146 | 0.109 | -0.282 | 0.778 |
| Crypturellus undulatus | precision0.5 | 1.183 | 0.049 | 1.088 | 1.278 | 24.369 | <.001 |
| Crypturellus undulatus | precision0.5:recall0.1 | 0.059 | 0.158 | -0.252 | 0.369 | 0.37 | 0.711 |
| Crypturellus undulatus | precision0.5:recall0.25 | 0.026 | 0.108 | -0.187 | 0.238 | 0.238 | 0.812 |
| Crypturellus undulatus | precision0.5:recall0.5 | -0.029 | 0.084 | -0.193 | 0.135 | -0.349 | 0.727 |
| Crypturellus undulatus | precision0.5:recall0.75 | -0.01 | 0.074 | -0.155 | 0.136 | -0.129 | 0.897 |
| Crypturellus undulatus | precision0.5:recall0.9 | -0.003 | 0.07 | -0.141 | 0.135 | -0.041 | 0.967 |
| Crypturellus undulatus | precision0.75 | 0.515 | 0.053 | 0.411 | 0.619 | 9.718 | <.001 |
| Crypturellus undulatus | precision0.75:recall0.1 | 0.06 | 0.173 | -0.28 | 0.4 | 0.346 | 0.729 |
| Crypturellus undulatus | precision0.75:recall0.25 | 0.054 | 0.118 | -0.178 | 0.286 | 0.457 | 0.647 |
| Crypturellus undulatus | precision0.75:recall0.5 | 0.007 | 0.091 | -0.172 | 0.186 | 0.081 | 0.936 |
| Crypturellus undulatus | precision0.75:recall0.75 | 0.023 | 0.081 | -0.135 | 0.182 | 0.287 | 0.774 |
| Crypturellus undulatus | precision0.75:recall0.9 | 0.021 | 0.077 | -0.129 | 0.172 | 0.276 | 0.783 |
| Crypturellus undulatus | precision0.9 | 0.21 | 0.056 | 0.1 | 0.319 | 3.758 | <.001 |
| Crypturellus undulatus | precision0.9:recall0.1 | 0.027 | 0.183 | -0.333 | 0.386 | 0.145 | 0.885 |
| Crypturellus undulatus | precision0.9:recall0.25 | 0.03 | 0.125 | -0.215 | 0.275 | 0.239 | 0.811 |
| Crypturellus undulatus | precision0.9:recall0.5 | 0.009 | 0.096 | -0.18 | 0.198 | 0.094 | 0.925 |
| Crypturellus undulatus | precision0.9:recall0.75 | -0.004 | 0.085 | -0.171 | 0.163 | -0.043 | 0.966 |
| Crypturellus undulatus | precision0.9:recall0.9 | 0.009 | 0.081 | -0.15 | 0.167 | 0.108 | 0.914 |
| Crypturellus undulatus | recall0.1 | -2.336 | 0.135 | -2.601 | -2.071 | -17.277 | <.001 |
| Crypturellus undulatus | recall0.25 | -1.462 | 0.092 | -1.642 | -1.281 | -15.877 | <.001 |
| Crypturellus undulatus | recall0.5 | -0.71 | 0.071 | -0.849 | -0.571 | -10.031 | <.001 |
| Crypturellus undulatus | recall0.75 | -0.305 | 0.063 | -0.428 | -0.183 | -4.881 | <.001 |
| Crypturellus undulatus | recall0.9 | -0.097 | 0.059 | -0.213 | 0.019 | -1.632 | 0.103 |
| Crypturellus undulatus | time | 3.296 | 0.171 | 2.962 | 3.631 | 19.293 | <.001 |
| Crypturellus undulatus | time:precision0.1 | -3.049 | 0.181 | -3.404 | -2.693 | -16.809 | <.001 |
| Crypturellus undulatus | time:precision0.1:recall0.1 | 0.103 | 0.584 | -1.042 | 1.249 | 0.176 | 0.86 |
| Crypturellus undulatus | time:precision0.1:recall0.25 | -0.164 | 0.399 | -0.946 | 0.618 | -0.411 | 0.681 |
| Crypturellus undulatus | time:precision0.1:recall0.5 | -0.014 | 0.31 | -0.622 | 0.593 | -0.046 | 0.963 |
| Crypturellus undulatus | time:precision0.1:recall0.75 | -0.103 | 0.275 | -0.643 | 0.437 | -0.374 | 0.708 |
| Crypturellus undulatus | time:precision0.1:recall0.9 | 0.013 | 0.263 | -0.502 | 0.528 | 0.05 | 0.96 |
| Crypturellus undulatus | time:precision0.25 | -2.611 | 0.19 | -2.983 | -2.239 | -13.765 | <.001 |
| Crypturellus undulatus | time:precision0.25:recall0.1 | -0.054 | 0.616 | -1.261 | 1.154 | -0.087 | 0.931 |
| Crypturellus undulatus | time:precision0.25:recall0.25 | -0.065 | 0.42 | -0.888 | 0.759 | -0.154 | 0.878 |
| Crypturellus undulatus | time:precision0.25:recall0.5 | 0.124 | 0.326 | -0.514 | 0.762 | 0.38 | 0.704 |
| Crypturellus undulatus | time:precision0.25:recall0.75 | -0.066 | 0.289 | -0.631 | 0.5 | -0.227 | 0.82 |
| Crypturellus undulatus | time:precision0.25:recall0.9 | 0.012 | 0.275 | -0.527 | 0.551 | 0.044 | 0.965 |
| Crypturellus undulatus | time:precision0.5 | -1.713 | 0.206 | -2.117 | -1.31 | -8.322 | <.001 |
| Crypturellus undulatus | time:precision0.5:recall0.1 | -0.477 | 0.669 | -1.789 | 0.834 | -0.713 | 0.476 |
| Crypturellus undulatus | time:precision0.5:recall0.25 | -0.257 | 0.457 | -1.153 | 0.638 | -0.564 | 0.573 |
| Crypturellus undulatus | time:precision0.5:recall0.5 | 0.042 | 0.354 | -0.652 | 0.736 | 0.118 | 0.906 |
| Crypturellus undulatus | time:precision0.5:recall0.75 | -0.062 | 0.314 | -0.677 | 0.553 | -0.197 | 0.843 |
| Crypturellus undulatus | time:precision0.5:recall0.9 | -0.02 | 0.298 | -0.605 | 0.565 | -0.067 | 0.946 |
| Crypturellus undulatus | time:precision0.75 | -0.838 | 0.223 | -1.276 | -0.4 | -3.752 | <.001 |
| Crypturellus undulatus | time:precision0.75:recall0.1 | -0.327 | 0.728 | -1.754 | 1.1 | -0.449 | 0.653 |
| Crypturellus undulatus | time:precision0.75:recall0.25 | -0.222 | 0.496 | -1.194 | 0.75 | -0.447 | 0.655 |
| Crypturellus undulatus | time:precision0.75:recall0.5 | -0.016 | 0.384 | -0.769 | 0.738 | -0.04 | 0.968 |
| Crypturellus undulatus | time:precision0.75:recall0.75 | -0.1 | 0.34 | -0.766 | 0.567 | -0.293 | 0.77 |
| Crypturellus undulatus | time:precision0.75:recall0.9 | -0.076 | 0.324 | -0.71 | 0.558 | -0.235 | 0.814 |
| Crypturellus undulatus | time:precision0.9 | -0.368 | 0.234 | -0.826 | 0.091 | -1.573 | 0.116 |
| Crypturellus undulatus | time:precision0.9:recall0.1 | -0.082 | 0.765 | -1.582 | 1.418 | -0.107 | 0.915 |
| Crypturellus undulatus | time:precision0.9:recall0.25 | -0.186 | 0.521 | -1.207 | 0.835 | -0.357 | 0.721 |
| Crypturellus undulatus | time:precision0.9:recall0.5 | -0.057 | 0.403 | -0.846 | 0.733 | -0.14 | 0.888 |
| Crypturellus undulatus | time:precision0.9:recall0.75 | -0.018 | 0.357 | -0.717 | 0.682 | -0.05 | 0.96 |
| Crypturellus undulatus | time:precision0.9:recall0.9 | -0.061 | 0.339 | -0.726 | 0.604 | -0.18 | 0.857 |
| Crypturellus undulatus | time:recall0.1 | -0.091 | 0.561 | -1.19 | 1.008 | -0.163 | 0.871 |
| Crypturellus undulatus | time:recall0.25 | 0.193 | 0.381 | -0.555 | 0.94 | 0.505 | 0.613 |
| Crypturellus undulatus | time:recall0.5 | -0.048 | 0.295 | -0.625 | 0.529 | -0.163 | 0.871 |
| Crypturellus undulatus | time:recall0.75 | 0.052 | 0.261 | -0.459 | 0.563 | 0.199 | 0.842 |
| Crypturellus undulatus | time:recall0.9 | -0.05 | 0.248 | -0.536 | 0.436 | -0.202 | 0.84 |
| Pygiptila stellaris | (Intercept) | -5.453 | 0.056 | -5.563 | -5.344 | -97.746 | <.001 |
| Pygiptila stellaris | I(time^2) | -16.233 | 0.275 | -16.773 | -15.693 | -58.955 | <.001 |
| Pygiptila stellaris | I(time^2):precision0.1 | 15.354 | 0.282 | 14.802 | 15.906 | 54.52 | <.001 |
| Pygiptila stellaris | I(time^2):precision0.1:recall0.1 | -0.845 | 0.882 | -2.574 | 0.883 | -0.958 | 0.338 |
| Pygiptila stellaris | I(time^2):precision0.1:recall0.25 | -0.867 | 0.603 | -2.048 | 0.315 | -1.438 | 0.151 |
| Pygiptila stellaris | I(time^2):precision0.1:recall0.5 | -0.336 | 0.48 | -1.276 | 0.604 | -0.701 | 0.483 |
| Pygiptila stellaris | I(time^2):precision0.1:recall0.75 | -0.017 | 0.428 | -0.856 | 0.822 | -0.039 | 0.969 |
| Pygiptila stellaris | I(time^2):precision0.1:recall0.9 | 0.057 | 0.409 | -0.744 | 0.859 | 0.14 | 0.889 |
| Pygiptila stellaris | I(time^2):precision0.25 | 13.734 | 0.287 | 13.17 | 14.297 | 47.779 | <.001 |
| Pygiptila stellaris | I(time^2):precision0.25:recall0.1 | -0.398 | 0.903 | -2.169 | 1.373 | -0.441 | 0.659 |
| Pygiptila stellaris | I(time^2):precision0.25:recall0.25 | -0.59 | 0.617 | -1.8 | 0.62 | -0.955 | 0.34 |
| Pygiptila stellaris | I(time^2):precision0.25:recall0.5 | -0.264 | 0.49 | -1.225 | 0.698 | -0.537 | 0.591 |
| Pygiptila stellaris | I(time^2):precision0.25:recall0.75 | -0.053 | 0.437 | -0.91 | 0.804 | -0.121 | 0.903 |
| Pygiptila stellaris | I(time^2):precision0.25:recall0.9 | 0.037 | 0.417 | -0.782 | 0.855 | 0.088 | 0.93 |
| Pygiptila stellaris | I(time^2):precision0.5 | 10.709 | 0.302 | 10.117 | 11.301 | 35.456 | <.001 |
| Pygiptila stellaris | I(time^2):precision0.5:recall0.1 | -0.35 | 0.952 | -2.215 | 1.515 | -0.368 | 0.713 |
| Pygiptila stellaris | I(time^2):precision0.5:recall0.25 | -0.491 | 0.65 | -1.765 | 0.784 | -0.755 | 0.45 |
| Pygiptila stellaris | I(time^2):precision0.5:recall0.5 | -0.107 | 0.516 | -1.118 | 0.904 | -0.208 | 0.835 |
| Pygiptila stellaris | I(time^2):precision0.5:recall0.75 | -0.039 | 0.46 | -0.94 | 0.861 | -0.086 | 0.932 |
| Pygiptila stellaris | I(time^2):precision0.5:recall0.9 | -0.011 | 0.439 | -0.871 | 0.848 | -0.026 | 0.979 |
| Pygiptila stellaris | I(time^2):precision0.75 | 6.634 | 0.328 | 5.991 | 7.277 | 20.223 | <.001 |
| Pygiptila stellaris | I(time^2):precision0.75:recall0.1 | -0.423 | 1.036 | -2.453 | 1.608 | -0.408 | 0.683 |
| Pygiptila stellaris | I(time^2):precision0.75:recall0.25 | -0.129 | 0.705 | -1.511 | 1.254 | -0.182 | 0.855 |
| Pygiptila stellaris | I(time^2):precision0.75:recall0.5 | -0.09 | 0.56 | -1.188 | 1.008 | -0.16 | 0.873 |
| Pygiptila stellaris | I(time^2):precision0.75:recall0.75 | 0.128 | 0.499 | -0.85 | 1.105 | 0.256 | 0.798 |
| Pygiptila stellaris | I(time^2):precision0.75:recall0.9 | -0.048 | 0.477 | -0.982 | 0.886 | -0.101 | 0.92 |
| Pygiptila stellaris | I(time^2):precision0.9 | 3.205 | 0.357 | 2.506 | 3.905 | 8.98 | <.001 |
| Pygiptila stellaris | I(time^2):precision0.9:recall0.1 | -0.717 | 1.135 | -2.941 | 1.507 | -0.632 | 0.528 |
| Pygiptila stellaris | I(time^2):precision0.9:recall0.25 | 0.032 | 0.766 | -1.47 | 1.534 | 0.042 | 0.967 |
| Pygiptila stellaris | I(time^2):precision0.9:recall0.5 | 0.043 | 0.609 | -1.151 | 1.237 | 0.07 | 0.944 |
| Pygiptila stellaris | I(time^2):precision0.9:recall0.75 | 0.109 | 0.543 | -0.954 | 1.173 | 0.202 | 0.84 |
| Pygiptila stellaris | I(time^2):precision0.9:recall0.9 | -0.124 | 0.519 | -1.142 | 0.893 | -0.24 | 0.811 |
| Pygiptila stellaris | I(time^2):recall0.1 | 1.387 | 0.868 | -0.313 | 3.088 | 1.599 | 0.11 |
| Pygiptila stellaris | I(time^2):recall0.25 | 1.145 | 0.592 | -0.016 | 2.305 | 1.933 | 0.053 |
| Pygiptila stellaris | I(time^2):recall0.5 | 0.509 | 0.47 | -0.413 | 1.431 | 1.082 | 0.279 |
| Pygiptila stellaris | I(time^2):recall0.75 | 0.159 | 0.419 | -0.663 | 0.981 | 0.379 | 0.704 |
| Pygiptila stellaris | I(time^2):recall0.9 | 0.001 | 0.4 | -0.783 | 0.785 | 0.002 | 0.999 |
| Pygiptila stellaris | precision0.1 | 4.955 | 0.057 | 4.843 | 5.068 | 86.419 | <.001 |
| Pygiptila stellaris | precision0.1:recall0.1 | -0.527 | 0.182 | -0.885 | -0.17 | -2.892 | 0.004 |
| Pygiptila stellaris | precision0.1:recall0.25 | -0.428 | 0.125 | -0.674 | -0.183 | -3.42 | <.001 |
| Pygiptila stellaris | precision0.1:recall0.5 | -0.284 | 0.098 | -0.477 | -0.091 | -2.886 | 0.004 |
| Pygiptila stellaris | precision0.1:recall0.75 | -0.15 | 0.087 | -0.321 | 0.021 | -1.723 | 0.085 |
| Pygiptila stellaris | precision0.1:recall0.9 | -0.035 | 0.083 | -0.198 | 0.128 | -0.421 | 0.674 |
| Pygiptila stellaris | precision0.25 | 3.488 | 0.059 | 3.373 | 3.603 | 59.336 | <.001 |
| Pygiptila stellaris | precision0.25:recall0.1 | -0.158 | 0.188 | -0.526 | 0.209 | -0.843 | 0.399 |
| Pygiptila stellaris | precision0.25:recall0.25 | -0.12 | 0.129 | -0.373 | 0.132 | -0.934 | 0.351 |
| Pygiptila stellaris | precision0.25:recall0.5 | -0.088 | 0.101 | -0.286 | 0.11 | -0.869 | 0.385 |
| Pygiptila stellaris | precision0.25:recall0.75 | -0.042 | 0.089 | -0.218 | 0.133 | -0.472 | 0.637 |
| Pygiptila stellaris | precision0.25:recall0.9 | 0.004 | 0.085 | -0.164 | 0.171 | 0.044 | 0.965 |
| Pygiptila stellaris | precision0.5 | 2.279 | 0.062 | 2.157 | 2.401 | 36.581 | <.001 |
| Pygiptila stellaris | precision0.5:recall0.1 | -0.069 | 0.199 | -0.46 | 0.321 | -0.347 | 0.728 |
| Pygiptila stellaris | precision0.5:recall0.25 | -0.031 | 0.137 | -0.299 | 0.236 | -0.23 | 0.818 |
| Pygiptila stellaris | precision0.5:recall0.5 | -0.003 | 0.107 | -0.213 | 0.207 | -0.032 | 0.975 |
| Pygiptila stellaris | precision0.5:recall0.75 | -0.026 | 0.095 | -0.212 | 0.16 | -0.276 | 0.782 |
| Pygiptila stellaris | precision0.5:recall0.9 | -0.005 | 0.091 | -0.182 | 0.173 | -0.052 | 0.959 |
| Pygiptila stellaris | precision0.75 | 1.274 | 0.068 | 1.141 | 1.407 | 18.794 | <.001 |
| Pygiptila stellaris | precision0.75:recall0.1 | -0.102 | 0.218 | -0.53 | 0.325 | -0.469 | 0.639 |
| Pygiptila stellaris | precision0.75:recall0.25 | 0.011 | 0.149 | -0.28 | 0.303 | 0.076 | 0.94 |
| Pygiptila stellaris | precision0.75:recall0.5 | -0.005 | 0.117 | -0.233 | 0.224 | -0.039 | 0.969 |
| Pygiptila stellaris | precision0.75:recall0.75 | -0.001 | 0.103 | -0.203 | 0.202 | -0.007 | 0.995 |
| Pygiptila stellaris | precision0.75:recall0.9 | -0.005 | 0.099 | -0.198 | 0.189 | -0.047 | 0.963 |
| Pygiptila stellaris | precision0.9 | 0.6 | 0.073 | 0.457 | 0.743 | 8.209 | <.001 |
| Pygiptila stellaris | precision0.9:recall0.1 | -0.168 | 0.237 | -0.634 | 0.297 | -0.709 | 0.479 |
| Pygiptila stellaris | precision0.9:recall0.25 | -0.002 | 0.161 | -0.316 | 0.313 | -0.01 | 0.992 |
| Pygiptila stellaris | precision0.9:recall0.5 | 0.013 | 0.126 | -0.233 | 0.26 | 0.107 | 0.915 |
| Pygiptila stellaris | precision0.9:recall0.75 | -0.008 | 0.111 | -0.227 | 0.21 | -0.076 | 0.94 |
| Pygiptila stellaris | precision0.9:recall0.9 | -0.037 | 0.107 | -0.246 | 0.172 | -0.346 | 0.729 |
| Pygiptila stellaris | recall0.1 | -2.174 | 0.179 | -2.524 | -1.823 | -12.158 | <.001 |
| Pygiptila stellaris | recall0.25 | -1.327 | 0.123 | -1.567 | -1.087 | -10.818 | <.001 |
| Pygiptila stellaris | recall0.5 | -0.681 | 0.096 | -0.87 | -0.493 | -7.086 | <.001 |
| Pygiptila stellaris | recall0.75 | -0.277 | 0.085 | -0.443 | -0.11 | -3.255 | 0.001 |
| Pygiptila stellaris | recall0.9 | -0.121 | 0.081 | -0.28 | 0.038 | -1.488 | 0.137 |
| Pygiptila stellaris | time | 14.279 | 0.255 | 13.778 | 14.78 | 55.898 | <.001 |
| Pygiptila stellaris | time:precision0.1 | -13.475 | 0.263 | -13.989 | -12.96 | -51.303 | <.001 |
| Pygiptila stellaris | time:precision0.1:recall0.1 | 0.622 | 0.828 | -1.001 | 2.244 | 0.751 | 0.453 |
| Pygiptila stellaris | time:precision0.1:recall0.25 | 0.565 | 0.567 | -0.546 | 1.677 | 0.997 | 0.319 |
| Pygiptila stellaris | time:precision0.1:recall0.5 | 0.209 | 0.449 | -0.671 | 1.088 | 0.465 | 0.642 |
| Pygiptila stellaris | time:precision0.1:recall0.75 | 0.015 | 0.399 | -0.767 | 0.797 | 0.038 | 0.97 |
| Pygiptila stellaris | time:precision0.1:recall0.9 | -0.088 | 0.381 | -0.835 | 0.66 | -0.23 | 0.818 |
| Pygiptila stellaris | time:precision0.25 | -12 | 0.269 | -12.527 | -11.473 | -44.603 | <.001 |
| Pygiptila stellaris | time:precision0.25:recall0.1 | 0.23 | 0.851 | -1.438 | 1.899 | 0.271 | 0.787 |
| Pygiptila stellaris | time:precision0.25:recall0.25 | 0.322 | 0.583 | -0.82 | 1.465 | 0.553 | 0.58 |
| Pygiptila stellaris | time:precision0.25:recall0.5 | 0.17 | 0.461 | -0.733 | 1.072 | 0.368 | 0.713 |
| Pygiptila stellaris | time:precision0.25:recall0.75 | 0.042 | 0.409 | -0.76 | 0.844 | 0.103 | 0.918 |
| Pygiptila stellaris | time:precision0.25:recall0.9 | -0.059 | 0.391 | -0.825 | 0.707 | -0.151 | 0.88 |
| Pygiptila stellaris | time:precision0.5 | -9.255 | 0.284 | -9.813 | -8.698 | -32.564 | <.001 |
| Pygiptila stellaris | time:precision0.5:recall0.1 | 0.21 | 0.902 | -1.558 | 1.978 | 0.233 | 0.816 |
| Pygiptila stellaris | time:precision0.5:recall0.25 | 0.234 | 0.617 | -0.976 | 1.443 | 0.378 | 0.705 |
| Pygiptila stellaris | time:precision0.5:recall0.5 | 0.004 | 0.487 | -0.951 | 0.958 | 0.007 | 0.994 |
| Pygiptila stellaris | time:precision0.5:recall0.75 | 0.054 | 0.433 | -0.794 | 0.901 | 0.124 | 0.902 |
| Pygiptila stellaris | time:precision0.5:recall0.9 | 0.009 | 0.413 | -0.8 | 0.819 | 0.022 | 0.982 |
| Pygiptila stellaris | time:precision0.75 | -5.686 | 0.309 | -6.291 | -5.081 | -18.421 | <.001 |
| Pygiptila stellaris | time:precision0.75:recall0.1 | 0.4 | 0.984 | -1.528 | 2.328 | 0.407 | 0.684 |
| Pygiptila stellaris | time:precision0.75:recall0.25 | -0.002 | 0.67 | -1.315 | 1.311 | -0.002 | 0.998 |
| Pygiptila stellaris | time:precision0.75:recall0.5 | 0.034 | 0.529 | -1.003 | 1.071 | 0.064 | 0.949 |
| Pygiptila stellaris | time:precision0.75:recall0.75 | -0.079 | 0.469 | -0.999 | 0.841 | -0.168 | 0.867 |
| Pygiptila stellaris | time:precision0.75:recall0.9 | 0.035 | 0.449 | -0.845 | 0.914 | 0.078 | 0.938 |
| Pygiptila stellaris | time:precision0.9 | -2.766 | 0.334 | -3.42 | -2.112 | -8.288 | <.001 |
| Pygiptila stellaris | time:precision0.9:recall0.1 | 0.742 | 1.071 | -1.358 | 2.841 | 0.693 | 0.489 |
| Pygiptila stellaris | time:precision0.9:recall0.25 | -0.035 | 0.724 | -1.454 | 1.385 | -0.048 | 0.962 |
| Pygiptila stellaris | time:precision0.9:recall0.5 | -0.063 | 0.572 | -1.184 | 1.057 | -0.111 | 0.912 |
| Pygiptila stellaris | time:precision0.9:recall0.75 | -0.041 | 0.508 | -1.036 | 0.954 | -0.081 | 0.936 |
| Pygiptila stellaris | time:precision0.9:recall0.9 | 0.147 | 0.486 | -0.805 | 1.099 | 0.302 | 0.762 |
| Pygiptila stellaris | time:recall0.1 | -1.146 | 0.811 | -2.736 | 0.444 | -1.412 | 0.158 |
| Pygiptila stellaris | time:recall0.25 | -0.838 | 0.555 | -1.926 | 0.249 | -1.511 | 0.131 |
| Pygiptila stellaris | time:recall0.5 | -0.367 | 0.438 | -1.225 | 0.492 | -0.838 | 0.402 |
| Pygiptila stellaris | time:recall0.75 | -0.144 | 0.389 | -0.907 | 0.618 | -0.371 | 0.711 |
| Pygiptila stellaris | time:recall0.9 | 0.02 | 0.371 | -0.708 | 0.748 | 0.053 | 0.957 |
| Micrastur ruficollis | (Intercept) | -4.255 | 0.046 | -4.344 | -4.165 | -93.245 | <.001 |
| Micrastur ruficollis | I(time^2) | -8.085 | 0.237 | -8.549 | -7.621 | -34.167 | <.001 |
| Micrastur ruficollis | I(time^2):precision0.1 | 7.477 | 0.245 | 6.996 | 7.958 | 30.48 | <.001 |
| Micrastur ruficollis | I(time^2):precision0.1:recall0.1 | 1.698 | 0.837 | 0.057 | 3.339 | 2.028 | 0.043 |
| Micrastur ruficollis | I(time^2):precision0.1:recall0.25 | 0.3 | 0.545 | -0.768 | 1.368 | 0.551 | 0.581 |
| Micrastur ruficollis | I(time^2):precision0.1:recall0.5 | -0.077 | 0.421 | -0.902 | 0.747 | -0.184 | 0.854 |
| Micrastur ruficollis | I(time^2):precision0.1:recall0.75 | -0.136 | 0.373 | -0.866 | 0.594 | -0.365 | 0.715 |
| Micrastur ruficollis | I(time^2):precision0.1:recall0.9 | -0.054 | 0.356 | -0.751 | 0.644 | -0.15 | 0.88 |
| Micrastur ruficollis | I(time^2):precision0.25 | 6.555 | 0.255 | 6.055 | 7.054 | 25.727 | <.001 |
| Micrastur ruficollis | I(time^2):precision0.25:recall0.1 | 1.379 | 0.869 | -0.324 | 3.083 | 1.587 | 0.113 |
| Micrastur ruficollis | I(time^2):precision0.25:recall0.25 | 0.414 | 0.567 | -0.697 | 1.524 | 0.73 | 0.465 |
| Micrastur ruficollis | I(time^2):precision0.25:recall0.5 | 0.029 | 0.438 | -0.829 | 0.886 | 0.066 | 0.948 |
| Micrastur ruficollis | I(time^2):precision0.25:recall0.75 | -0.144 | 0.387 | -0.903 | 0.615 | -0.371 | 0.71 |
| Micrastur ruficollis | I(time^2):precision0.25:recall0.9 | -0.007 | 0.37 | -0.731 | 0.718 | -0.018 | 0.986 |
| Micrastur ruficollis | I(time^2):precision0.5 | 4.861 | 0.273 | 4.325 | 5.397 | 17.778 | <.001 |
| Micrastur ruficollis | I(time^2):precision0.5:recall0.1 | 1.258 | 0.93 | -0.565 | 3.081 | 1.352 | 0.176 |
| Micrastur ruficollis | I(time^2):precision0.5:recall0.25 | 0.161 | 0.609 | -1.033 | 1.355 | 0.265 | 0.791 |
| Micrastur ruficollis | I(time^2):precision0.5:recall0.5 | 0.214 | 0.47 | -0.706 | 1.135 | 0.456 | 0.648 |
| Micrastur ruficollis | I(time^2):precision0.5:recall0.75 | 0.029 | 0.416 | -0.786 | 0.843 | 0.069 | 0.945 |
| Micrastur ruficollis | I(time^2):precision0.5:recall0.9 | 0.112 | 0.397 | -0.665 | 0.89 | 0.283 | 0.777 |
| Micrastur ruficollis | I(time^2):precision0.75 | 2.841 | 0.298 | 2.257 | 3.424 | 9.541 | <.001 |
| Micrastur ruficollis | I(time^2):precision0.75:recall0.1 | 0.73 | 1.015 | -1.26 | 2.719 | 0.719 | 0.472 |
| Micrastur ruficollis | I(time^2):precision0.75:recall0.25 | -0.238 | 0.667 | -1.545 | 1.068 | -0.357 | 0.721 |
| Micrastur ruficollis | I(time^2):precision0.75:recall0.5 | 0.119 | 0.511 | -0.883 | 1.122 | 0.233 | 0.816 |
| Micrastur ruficollis | I(time^2):precision0.75:recall0.75 | -0.012 | 0.453 | -0.899 | 0.876 | -0.026 | 0.979 |
| Micrastur ruficollis | I(time^2):precision0.75:recall0.9 | 0.105 | 0.432 | -0.742 | 0.951 | 0.243 | 0.808 |
| Micrastur ruficollis | I(time^2):precision0.9 | 1.209 | 0.318 | 0.586 | 1.832 | 3.801 | <.001 |
| Micrastur ruficollis | I(time^2):precision0.9:recall0.1 | 0.463 | 1.087 | -1.668 | 2.593 | 0.426 | 0.67 |
| Micrastur ruficollis | I(time^2):precision0.9:recall0.25 | 0.035 | 0.709 | -1.355 | 1.426 | 0.05 | 0.96 |
| Micrastur ruficollis | I(time^2):precision0.9:recall0.5 | -0.064 | 0.548 | -1.137 | 1.01 | -0.116 | 0.908 |
| Micrastur ruficollis | I(time^2):precision0.9:recall0.75 | 0.063 | 0.483 | -0.884 | 1.01 | 0.131 | 0.896 |
| Micrastur ruficollis | I(time^2):precision0.9:recall0.9 | 0.168 | 0.461 | -0.736 | 1.071 | 0.363 | 0.716 |
| Micrastur ruficollis | I(time^2):recall0.1 | -1.248 | 0.816 | -2.848 | 0.352 | -1.529 | 0.126 |
| Micrastur ruficollis | I(time^2):recall0.25 | 0.014 | 0.529 | -1.023 | 1.051 | 0.027 | 0.979 |
| Micrastur ruficollis | I(time^2):recall0.5 | 0.202 | 0.407 | -0.596 | 1.001 | 0.497 | 0.619 |
| Micrastur ruficollis | I(time^2):recall0.75 | 0.184 | 0.36 | -0.522 | 0.889 | 0.511 | 0.61 |
| Micrastur ruficollis | I(time^2):recall0.9 | 0.031 | 0.344 | -0.642 | 0.704 | 0.09 | 0.928 |
| Micrastur ruficollis | precision0.1 | 3.268 | 0.048 | 3.174 | 3.362 | 68.291 | <.001 |
| Micrastur ruficollis | precision0.1:recall0.1 | 0.053 | 0.165 | -0.271 | 0.377 | 0.321 | 0.748 |
| Micrastur ruficollis | precision0.1:recall0.25 | -0.241 | 0.105 | -0.447 | -0.035 | -2.295 | 0.022 |
| Micrastur ruficollis | precision0.1:recall0.5 | -0.19 | 0.082 | -0.35 | -0.03 | -2.321 | 0.02 |
| Micrastur ruficollis | precision0.1:recall0.75 | -0.128 | 0.072 | -0.27 | 0.014 | -1.763 | 0.078 |
| Micrastur ruficollis | precision0.1:recall0.9 | -0.049 | 0.069 | -0.185 | 0.087 | -0.703 | 0.482 |
| Micrastur ruficollis | precision0.25 | 2.058 | 0.05 | 1.959 | 2.156 | 41.01 | <.001 |
| Micrastur ruficollis | precision0.25:recall0.1 | 0.199 | 0.173 | -0.141 | 0.538 | 1.148 | 0.251 |
| Micrastur ruficollis | precision0.25:recall0.25 | -0.026 | 0.11 | -0.242 | 0.19 | -0.236 | 0.814 |
| Micrastur ruficollis | precision0.25:recall0.5 | -0.072 | 0.086 | -0.24 | 0.096 | -0.837 | 0.402 |
| Micrastur ruficollis | precision0.25:recall0.75 | -0.07 | 0.076 | -0.219 | 0.079 | -0.922 | 0.356 |
| Micrastur ruficollis | precision0.25:recall0.9 | -0.011 | 0.073 | -0.153 | 0.132 | -0.146 | 0.884 |
| Micrastur ruficollis | precision0.5 | 1.138 | 0.054 | 1.031 | 1.245 | 20.915 | <.001 |
| Micrastur ruficollis | precision0.5:recall0.1 | 0.204 | 0.187 | -0.163 | 0.571 | 1.089 | 0.276 |
| Micrastur ruficollis | precision0.5:recall0.25 | 0.021 | 0.12 | -0.214 | 0.255 | 0.173 | 0.863 |
| Micrastur ruficollis | precision0.5:recall0.5 | 0.034 | 0.093 | -0.148 | 0.216 | 0.367 | 0.714 |
| Micrastur ruficollis | precision0.5:recall0.75 | 0.018 | 0.082 | -0.144 | 0.179 | 0.212 | 0.832 |
| Micrastur ruficollis | precision0.5:recall0.9 | 0.029 | 0.079 | -0.126 | 0.184 | 0.366 | 0.714 |
| Micrastur ruficollis | precision0.75 | 0.574 | 0.059 | 0.459 | 0.69 | 9.774 | <.001 |
| Micrastur ruficollis | precision0.75:recall0.1 | 0.104 | 0.204 | -0.295 | 0.504 | 0.51 | 0.61 |
| Micrastur ruficollis | precision0.75:recall0.25 | -0.023 | 0.13 | -0.278 | 0.232 | -0.178 | 0.859 |
| Micrastur ruficollis | precision0.75:recall0.5 | -0.008 | 0.101 | -0.205 | 0.19 | -0.077 | 0.938 |
| Micrastur ruficollis | precision0.75:recall0.75 | -0.018 | 0.089 | -0.193 | 0.156 | -0.205 | 0.837 |
| Micrastur ruficollis | precision0.75:recall0.9 | 0.025 | 0.085 | -0.142 | 0.192 | 0.297 | 0.767 |
| Micrastur ruficollis | precision0.9 | 0.241 | 0.062 | 0.12 | 0.363 | 3.894 | <.001 |
| Micrastur ruficollis | precision0.9:recall0.1 | 0.082 | 0.215 | -0.34 | 0.503 | 0.379 | 0.705 |
| Micrastur ruficollis | precision0.9:recall0.25 | -0.034 | 0.137 | -0.303 | 0.235 | -0.246 | 0.806 |
| Micrastur ruficollis | precision0.9:recall0.5 | -0.016 | 0.106 | -0.225 | 0.192 | -0.151 | 0.88 |
| Micrastur ruficollis | precision0.9:recall0.75 | -0.012 | 0.094 | -0.196 | 0.172 | -0.129 | 0.897 |
| Micrastur ruficollis | precision0.9:recall0.9 | 0.013 | 0.09 | -0.163 | 0.189 | 0.147 | 0.883 |
| Micrastur ruficollis | recall0.1 | -2.594 | 0.16 | -2.908 | -2.28 | -16.203 | <.001 |
| Micrastur ruficollis | recall0.25 | -1.378 | 0.101 | -1.575 | -1.18 | -13.651 | <.001 |
| Micrastur ruficollis | recall0.5 | -0.669 | 0.078 | -0.822 | -0.515 | -8.54 | <.001 |
| Micrastur ruficollis | recall0.75 | -0.253 | 0.069 | -0.388 | -0.117 | -3.65 | <.001 |
| Micrastur ruficollis | recall0.9 | -0.105 | 0.066 | -0.235 | 0.025 | -1.589 | 0.112 |
| Micrastur ruficollis | time | 6.382 | 0.219 | 5.952 | 6.812 | 29.098 | <.001 |
| Micrastur ruficollis | time:precision0.1 | -5.879 | 0.229 | -6.329 | -5.43 | -25.649 | <.001 |
| Micrastur ruficollis | time:precision0.1:recall0.1 | -1.698 | 0.782 | -3.23 | -0.166 | -2.172 | 0.03 |
| Micrastur ruficollis | time:precision0.1:recall0.25 | -0.165 | 0.506 | -1.157 | 0.826 | -0.327 | 0.744 |
| Micrastur ruficollis | time:precision0.1:recall0.5 | 0.099 | 0.392 | -0.67 | 0.868 | 0.253 | 0.801 |
| Micrastur ruficollis | time:precision0.1:recall0.75 | 0.17 | 0.348 | -0.512 | 0.851 | 0.488 | 0.625 |
| Micrastur ruficollis | time:precision0.1:recall0.9 | 0.061 | 0.333 | -0.591 | 0.713 | 0.183 | 0.855 |
| Micrastur ruficollis | time:precision0.25 | -5.111 | 0.24 | -5.58 | -4.641 | -21.335 | <.001 |
| Micrastur ruficollis | time:precision0.25:recall0.1 | -1.391 | 0.817 | -2.991 | 0.21 | -1.703 | 0.089 |
| Micrastur ruficollis | time:precision0.25:recall0.25 | -0.336 | 0.529 | -1.373 | 0.702 | -0.634 | 0.526 |
| Micrastur ruficollis | time:precision0.25:recall0.5 | 0.03 | 0.411 | -0.775 | 0.836 | 0.074 | 0.941 |
| Micrastur ruficollis | time:precision0.25:recall0.75 | 0.185 | 0.364 | -0.528 | 0.898 | 0.508 | 0.612 |
| Micrastur ruficollis | time:precision0.25:recall0.9 | 0.005 | 0.348 | -0.676 | 0.687 | 0.015 | 0.988 |
| Micrastur ruficollis | time:precision0.5 | -3.705 | 0.259 | -4.212 | -3.199 | -14.326 | <.001 |
| Micrastur ruficollis | time:precision0.5:recall0.1 | -1.209 | 0.88 | -2.933 | 0.516 | -1.374 | 0.169 |
| Micrastur ruficollis | time:precision0.5:recall0.25 | -0.177 | 0.572 | -1.299 | 0.945 | -0.309 | 0.758 |
| Micrastur ruficollis | time:precision0.5:recall0.5 | -0.238 | 0.443 | -1.107 | 0.631 | -0.537 | 0.591 |
| Micrastur ruficollis | time:precision0.5:recall0.75 | -0.073 | 0.393 | -0.843 | 0.696 | -0.187 | 0.852 |
| Micrastur ruficollis | time:precision0.5:recall0.9 | -0.138 | 0.375 | -0.873 | 0.598 | -0.367 | 0.713 |
| Micrastur ruficollis | time:precision0.75 | -2.207 | 0.28 | -2.756 | -1.657 | -7.872 | <.001 |
| Micrastur ruficollis | time:precision0.75:recall0.1 | -0.652 | 0.958 | -2.529 | 1.225 | -0.681 | 0.496 |
| Micrastur ruficollis | time:precision0.75:recall0.25 | 0.181 | 0.623 | -1.041 | 1.402 | 0.29 | 0.772 |
| Micrastur ruficollis | time:precision0.75:recall0.5 | -0.065 | 0.481 | -1.009 | 0.878 | -0.136 | 0.892 |
| Micrastur ruficollis | time:precision0.75:recall0.75 | 0.041 | 0.426 | -0.794 | 0.876 | 0.095 | 0.924 |
| Micrastur ruficollis | time:precision0.75:recall0.9 | -0.122 | 0.407 | -0.919 | 0.675 | -0.3 | 0.764 |
| Micrastur ruficollis | time:precision0.9 | -0.963 | 0.297 | -1.545 | -0.381 | -3.242 | 0.001 |
| Micrastur ruficollis | time:precision0.9:recall0.1 | -0.43 | 1.016 | -2.422 | 1.562 | -0.423 | 0.672 |
| Micrastur ruficollis | time:precision0.9:recall0.25 | 0.057 | 0.66 | -1.236 | 1.351 | 0.087 | 0.931 |
| Micrastur ruficollis | time:precision0.9:recall0.5 | 0.068 | 0.511 | -0.933 | 1.069 | 0.133 | 0.894 |
| Micrastur ruficollis | time:precision0.9:recall0.75 | -0.009 | 0.451 | -0.893 | 0.875 | -0.019 | 0.985 |
| Micrastur ruficollis | time:precision0.9:recall0.9 | -0.124 | 0.431 | -0.968 | 0.719 | -0.289 | 0.773 |
| Micrastur ruficollis | time:recall0.1 | 1.278 | 0.758 | -0.207 | 2.763 | 1.687 | 0.092 |
| Micrastur ruficollis | time:recall0.25 | -0.094 | 0.487 | -1.049 | 0.862 | -0.193 | 0.847 |
| Micrastur ruficollis | time:recall0.5 | -0.21 | 0.377 | -0.949 | 0.529 | -0.556 | 0.578 |
| Micrastur ruficollis | time:recall0.75 | -0.204 | 0.333 | -0.857 | 0.45 | -0.611 | 0.541 |
| Micrastur ruficollis | time:recall0.9 | -0.026 | 0.318 | -0.65 | 0.598 | -0.083 | 0.934 |
| Formicarius analis | (Intercept) | -4.547 | 0.038 | -4.621 | -4.473 | -120.235 | <.001 |
| Formicarius analis | I(time^2) | -5.401 | 0.114 | -5.625 | -5.177 | -47.261 | <.001 |
| Formicarius analis | I(time^2):precision0.25 | 4.776 | 0.128 | 4.525 | 5.027 | 37.26 | <.001 |
| Formicarius analis | I(time^2):precision0.25:recall0.1 | 0.393 | 0.397 | -0.385 | 1.171 | 0.989 | 0.322 |
| Formicarius analis | I(time^2):precision0.25:recall0.25 | -0.075 | 0.269 | -0.602 | 0.452 | -0.28 | 0.78 |
| Formicarius analis | I(time^2):precision0.25:recall0.5 | -0.005 | 0.213 | -0.422 | 0.412 | -0.023 | 0.982 |
| Formicarius analis | I(time^2):precision0.25:recall0.75 | 0.012 | 0.192 | -0.364 | 0.389 | 0.064 | 0.949 |
| Formicarius analis | I(time^2):precision0.25:recall0.9 | 0.039 | 0.185 | -0.324 | 0.401 | 0.209 | 0.835 |
| Formicarius analis | I(time^2):precision0.5 | 3.823 | 0.134 | 3.56 | 4.086 | 28.473 | <.001 |
| Formicarius analis | I(time^2):precision0.5:recall0.1 | 0.455 | 0.424 | -0.376 | 1.285 | 1.073 | 0.283 |
| Formicarius analis | I(time^2):precision0.5:recall0.25 | -0.083 | 0.286 | -0.645 | 0.478 | -0.291 | 0.771 |
| Formicarius analis | I(time^2):precision0.5:recall0.5 | -0.017 | 0.225 | -0.459 | 0.424 | -0.077 | 0.938 |
| Formicarius analis | I(time^2):precision0.5:recall0.75 | -0.039 | 0.202 | -0.436 | 0.358 | -0.193 | 0.847 |
| Formicarius analis | I(time^2):precision0.5:recall0.9 | -0.028 | 0.194 | -0.408 | 0.353 | -0.143 | 0.887 |
| Formicarius analis | I(time^2):precision0.75 | 2.395 | 0.144 | 2.112 | 2.678 | 16.589 | <.001 |
| Formicarius analis | I(time^2):precision0.75:recall0.1 | 0.427 | 0.458 | -0.472 | 1.325 | 0.931 | 0.352 |
| Formicarius analis | I(time^2):precision0.75:recall0.25 | -0.025 | 0.31 | -0.633 | 0.582 | -0.081 | 0.935 |
| Formicarius analis | I(time^2):precision0.75:recall0.5 | 0.082 | 0.243 | -0.394 | 0.558 | 0.339 | 0.735 |
| Formicarius analis | I(time^2):precision0.75:recall0.75 | 0.029 | 0.218 | -0.398 | 0.456 | 0.132 | 0.895 |
| Formicarius analis | I(time^2):precision0.75:recall0.9 | -0.019 | 0.209 | -0.429 | 0.39 | -0.093 | 0.926 |
| Formicarius analis | I(time^2):precision0.9 | 1.087 | 0.153 | 0.787 | 1.388 | 7.089 | <.001 |
| Formicarius analis | I(time^2):precision0.9:recall0.1 | 0.376 | 0.489 | -0.584 | 1.335 | 0.768 | 0.443 |
| Formicarius analis | I(time^2):precision0.9:recall0.25 | -0.046 | 0.331 | -0.695 | 0.602 | -0.139 | 0.889 |
| Formicarius analis | I(time^2):precision0.9:recall0.5 | 0.029 | 0.259 | -0.478 | 0.536 | 0.111 | 0.911 |
| Formicarius analis | I(time^2):precision0.9:recall0.75 | 0.081 | 0.232 | -0.373 | 0.535 | 0.35 | 0.726 |
| Formicarius analis | I(time^2):precision0.9:recall0.9 | 0.082 | 0.222 | -0.353 | 0.516 | 0.368 | 0.713 |
| Formicarius analis | I(time^2):recall0.1 | -0.21 | 0.368 | -0.931 | 0.512 | -0.569 | 0.569 |
| Formicarius analis | I(time^2):recall0.25 | 0.34 | 0.247 | -0.143 | 0.824 | 1.379 | 0.168 |
| Formicarius analis | I(time^2):recall0.5 | 0.332 | 0.193 | -0.047 | 0.71 | 1.719 | 0.086 |
| Formicarius analis | I(time^2):recall0.75 | 0.137 | 0.173 | -0.201 | 0.476 | 0.795 | 0.427 |
| Formicarius analis | I(time^2):recall0.9 | 0.066 | 0.165 | -0.258 | 0.39 | 0.399 | 0.69 |
| Formicarius analis | precision0.25 | 4.125 | 0.04 | 4.046 | 4.203 | 103.064 | <.001 |
| Formicarius analis | precision0.25:recall0.1 | -0.342 | 0.13 | -0.597 | -0.086 | -2.623 | 0.009 |
| Formicarius analis | precision0.25:recall0.25 | -0.406 | 0.087 | -0.577 | -0.236 | -4.667 | <.001 |
| Formicarius analis | precision0.25:recall0.5 | -0.33 | 0.067 | -0.462 | -0.197 | -4.884 | <.001 |
| Formicarius analis | precision0.25:recall0.75 | -0.189 | 0.06 | -0.307 | -0.071 | -3.128 | 0.002 |
| Formicarius analis | precision0.25:recall0.9 | -0.08 | 0.058 | -0.193 | 0.034 | -1.376 | 0.169 |
| Formicarius analis | precision0.5 | 2.568 | 0.042 | 2.486 | 2.65 | 61.465 | <.001 |
| Formicarius analis | precision0.5:recall0.1 | 0.023 | 0.137 | -0.245 | 0.292 | 0.172 | 0.864 |
| Formicarius analis | precision0.5:recall0.25 | -0.086 | 0.091 | -0.265 | 0.094 | -0.936 | 0.349 |
| Formicarius analis | precision0.5:recall0.5 | -0.081 | 0.071 | -0.22 | 0.057 | -1.149 | 0.251 |
| Formicarius analis | precision0.5:recall0.75 | -0.06 | 0.063 | -0.184 | 0.064 | -0.946 | 0.344 |
| Formicarius analis | precision0.5:recall0.9 | -0.026 | 0.06 | -0.144 | 0.093 | -0.423 | 0.673 |
| Formicarius analis | precision0.75 | 1.402 | 0.045 | 1.313 | 1.491 | 30.852 | <.001 |
| Formicarius analis | precision0.75:recall0.1 | 0.125 | 0.149 | -0.167 | 0.416 | 0.839 | 0.401 |
| Formicarius analis | precision0.75:recall0.25 | -0.019 | 0.1 | -0.214 | 0.177 | -0.187 | 0.852 |
| Formicarius analis | precision0.75:recall0.5 | 0.014 | 0.077 | -0.137 | 0.165 | 0.185 | 0.853 |
| Formicarius analis | precision0.75:recall0.75 | 0.011 | 0.069 | -0.124 | 0.145 | 0.158 | 0.875 |
| Formicarius analis | precision0.75:recall0.9 | -0.012 | 0.066 | -0.141 | 0.117 | -0.182 | 0.856 |
| Formicarius analis | precision0.9 | 0.611 | 0.049 | 0.514 | 0.708 | 12.357 | <.001 |
| Formicarius analis | precision0.9:recall0.1 | 0.104 | 0.162 | -0.214 | 0.423 | 0.643 | 0.52 |
| Formicarius analis | precision0.9:recall0.25 | -0.031 | 0.109 | -0.245 | 0.183 | -0.287 | 0.774 |
| Formicarius analis | precision0.9:recall0.5 | 0.001 | 0.084 | -0.163 | 0.166 | 0.015 | 0.988 |
| Formicarius analis | precision0.9:recall0.75 | 0.028 | 0.075 | -0.118 | 0.175 | 0.379 | 0.705 |
| Formicarius analis | precision0.9:recall0.9 | 0.024 | 0.072 | -0.116 | 0.164 | 0.332 | 0.74 |
| Formicarius analis | recall0.1 | -2.415 | 0.126 | -2.661 | -2.169 | -19.216 | <.001 |
| Formicarius analis | recall0.25 | -1.376 | 0.083 | -1.54 | -1.213 | -16.495 | <.001 |
| Formicarius analis | recall0.5 | -0.62 | 0.064 | -0.746 | -0.494 | -9.646 | <.001 |
| Formicarius analis | recall0.75 | -0.254 | 0.057 | -0.366 | -0.141 | -4.426 | <.001 |
| Formicarius analis | recall0.9 | -0.09 | 0.055 | -0.198 | 0.017 | -1.651 | 0.099 |
| Formicarius analis | time | 8.211 | 0.136 | 7.944 | 8.478 | 60.289 | <.001 |
| Formicarius analis | time:precision0.25 | -7.103 | 0.149 | -7.395 | -6.811 | -47.707 | <.001 |
| Formicarius analis | time:precision0.25:recall0.1 | -0.516 | 0.47 | -1.437 | 0.404 | -1.099 | 0.272 |
| Formicarius analis | time:precision0.25:recall0.25 | 0.019 | 0.317 | -0.602 | 0.639 | 0.059 | 0.953 |
| Formicarius analis | time:precision0.25:recall0.5 | 0.02 | 0.249 | -0.467 | 0.508 | 0.082 | 0.935 |
| Formicarius analis | time:precision0.25:recall0.75 | 0.012 | 0.224 | -0.426 | 0.45 | 0.054 | 0.957 |
| Formicarius analis | time:precision0.25:recall0.9 | -0.02 | 0.215 | -0.441 | 0.401 | -0.093 | 0.926 |
| Formicarius analis | time:precision0.5 | -5.558 | 0.156 | -5.864 | -5.253 | -35.659 | <.001 |
| Formicarius analis | time:precision0.5:recall0.1 | -0.575 | 0.498 | -1.552 | 0.402 | -1.154 | 0.249 |
| Formicarius analis | time:precision0.5:recall0.25 | 0.013 | 0.336 | -0.645 | 0.671 | 0.04 | 0.968 |
| Formicarius analis | time:precision0.5:recall0.5 | 0.006 | 0.262 | -0.509 | 0.52 | 0.021 | 0.983 |
| Formicarius analis | time:precision0.5:recall0.75 | 0.054 | 0.235 | -0.407 | 0.515 | 0.228 | 0.82 |
| Formicarius analis | time:precision0.5:recall0.9 | 0.032 | 0.225 | -0.41 | 0.474 | 0.141 | 0.888 |
| Formicarius analis | time:precision0.75 | -3.425 | 0.169 | -3.755 | -3.095 | -20.323 | <.001 |
| Formicarius analis | time:precision0.75:recall0.1 | -0.567 | 0.541 | -1.627 | 0.492 | -1.049 | 0.294 |
| Formicarius analis | time:precision0.75:recall0.25 | -0.006 | 0.365 | -0.721 | 0.709 | -0.016 | 0.987 |
| Formicarius analis | time:precision0.75:recall0.5 | -0.127 | 0.284 | -0.684 | 0.43 | -0.447 | 0.655 |
| Formicarius analis | time:precision0.75:recall0.75 | -0.063 | 0.254 | -0.561 | 0.436 | -0.246 | 0.806 |
| Formicarius analis | time:precision0.75:recall0.9 | 0.024 | 0.244 | -0.454 | 0.502 | 0.098 | 0.922 |
| Formicarius analis | time:precision0.9 | -1.554 | 0.181 | -1.909 | -1.2 | -8.594 | <.001 |
| Formicarius analis | time:precision0.9:recall0.1 | -0.453 | 0.583 | -1.595 | 0.689 | -0.777 | 0.437 |
| Formicarius analis | time:precision0.9:recall0.25 | 0.065 | 0.393 | -0.706 | 0.836 | 0.166 | 0.868 |
| Formicarius analis | time:precision0.9:recall0.5 | -0.037 | 0.306 | -0.636 | 0.562 | -0.122 | 0.903 |
| Formicarius analis | time:precision0.9:recall0.75 | -0.112 | 0.273 | -0.647 | 0.423 | -0.409 | 0.683 |
| Formicarius analis | time:precision0.9:recall0.9 | -0.1 | 0.261 | -0.612 | 0.413 | -0.382 | 0.703 |
| Formicarius analis | time:recall0.1 | 0.087 | 0.444 | -0.782 | 0.957 | 0.197 | 0.844 |
| Formicarius analis | time:recall0.25 | -0.462 | 0.296 | -1.043 | 0.119 | -1.559 | 0.119 |
| Formicarius analis | time:recall0.5 | -0.491 | 0.231 | -0.943 | -0.039 | -2.131 | 0.033 |
| Formicarius analis | time:recall0.75 | -0.223 | 0.206 | -0.627 | 0.181 | -1.084 | 0.279 |
| Formicarius analis | time:recall0.9 | -0.1 | 0.197 | -0.486 | 0.287 | -0.505 | 0.613 |
| Lipaugus vociferans | (Intercept) | -7.359 | 0.099 | -7.553 | -7.166 | -74.547 | <.001 |
| Lipaugus vociferans | I(time^2) | -3.875 | 0.211 | -4.289 | -3.461 | -18.346 | <.001 |
| Lipaugus vociferans | I(time^2):precision0.1 | 4.472 | 0.221 | 4.039 | 4.905 | 20.24 | <.001 |
| Lipaugus vociferans | I(time^2):precision0.1:recall0.1 | 0.001 | 0.69 | -1.351 | 1.354 | 0.002 | 0.998 |
| Lipaugus vociferans | I(time^2):precision0.1:recall0.25 | 0.019 | 0.47 | -0.902 | 0.941 | 0.041 | 0.967 |
| Lipaugus vociferans | I(time^2):precision0.1:recall0.5 | 0.045 | 0.373 | -0.686 | 0.777 | 0.122 | 0.903 |
| Lipaugus vociferans | I(time^2):precision0.1:recall0.75 | 0.025 | 0.334 | -0.63 | 0.68 | 0.075 | 0.94 |
| Lipaugus vociferans | I(time^2):precision0.1:recall0.9 | -0.121 | 0.319 | -0.746 | 0.505 | -0.378 | 0.706 |
| Lipaugus vociferans | I(time^2):precision0.25 | 5.08 | 0.221 | 4.648 | 5.513 | 23.019 | <.001 |
| Lipaugus vociferans | I(time^2):precision0.25:recall0.1 | 0.087 | 0.702 | -1.289 | 1.464 | 0.124 | 0.901 |
| Lipaugus vociferans | I(time^2):precision0.25:recall0.25 | 0.049 | 0.477 | -0.886 | 0.984 | 0.103 | 0.918 |
| Lipaugus vociferans | I(time^2):precision0.25:recall0.5 | 0.119 | 0.377 | -0.621 | 0.858 | 0.314 | 0.753 |
| Lipaugus vociferans | I(time^2):precision0.25:recall0.75 | 0.19 | 0.336 | -0.469 | 0.849 | 0.565 | 0.572 |
| Lipaugus vociferans | I(time^2):precision0.25:recall0.9 | 0.037 | 0.32 | -0.591 | 0.664 | 0.114 | 0.909 |
| Lipaugus vociferans | I(time^2):precision0.5 | 5.505 | 0.228 | 5.058 | 5.952 | 24.157 | <.001 |
| Lipaugus vociferans | I(time^2):precision0.5:recall0.1 | 0.267 | 0.726 | -1.157 | 1.69 | 0.367 | 0.713 |
| Lipaugus vociferans | I(time^2):precision0.5:recall0.25 | 0.094 | 0.494 | -0.874 | 1.063 | 0.191 | 0.848 |
| Lipaugus vociferans | I(time^2):precision0.5:recall0.5 | 0.153 | 0.39 | -0.612 | 0.918 | 0.392 | 0.695 |
| Lipaugus vociferans | I(time^2):precision0.5:recall0.75 | 0.213 | 0.348 | -0.468 | 0.894 | 0.614 | 0.539 |
| Lipaugus vociferans | I(time^2):precision0.5:recall0.9 | 0.045 | 0.331 | -0.603 | 0.692 | 0.135 | 0.893 |
| Lipaugus vociferans | I(time^2):precision0.75 | 5.008 | 0.24 | 4.538 | 5.478 | 20.867 | <.001 |
| Lipaugus vociferans | I(time^2):precision0.75:recall0.1 | 0.102 | 0.766 | -1.4 | 1.603 | 0.133 | 0.895 |
| Lipaugus vociferans | I(time^2):precision0.75:recall0.25 | -0.31 | 0.523 | -1.334 | 0.714 | -0.593 | 0.553 |
| Lipaugus vociferans | I(time^2):precision0.75:recall0.5 | -0.023 | 0.412 | -0.83 | 0.784 | -0.055 | 0.956 |
| Lipaugus vociferans | I(time^2):precision0.75:recall0.75 | 0.086 | 0.366 | -0.633 | 0.804 | 0.233 | 0.815 |
| Lipaugus vociferans | I(time^2):precision0.75:recall0.9 | 0.027 | 0.348 | -0.656 | 0.709 | 0.076 | 0.939 |
| Lipaugus vociferans | I(time^2):precision0.9 | 3.3 | 0.259 | 2.793 | 3.807 | 12.766 | <.001 |
| Lipaugus vociferans | I(time^2):precision0.9:recall0.1 | 0.241 | 0.824 | -1.375 | 1.856 | 0.292 | 0.77 |
| Lipaugus vociferans | I(time^2):precision0.9:recall0.25 | -0.193 | 0.564 | -1.297 | 0.912 | -0.342 | 0.733 |
| Lipaugus vociferans | I(time^2):precision0.9:recall0.5 | -0.046 | 0.444 | -0.917 | 0.824 | -0.104 | 0.917 |
| Lipaugus vociferans | I(time^2):precision0.9:recall0.75 | -0.01 | 0.395 | -0.784 | 0.764 | -0.026 | 0.98 |
| Lipaugus vociferans | I(time^2):precision0.9:recall0.9 | 0.077 | 0.375 | -0.657 | 0.812 | 0.207 | 0.836 |
| Lipaugus vociferans | I(time^2):recall0.1 | -0.377 | 0.678 | -1.706 | 0.952 | -0.557 | 0.578 |
| Lipaugus vociferans | I(time^2):recall0.25 | -0.461 | 0.459 | -1.362 | 0.439 | -1.004 | 0.315 |
| Lipaugus vociferans | I(time^2):recall0.5 | -0.447 | 0.363 | -1.158 | 0.264 | -1.232 | 0.218 |
| Lipaugus vociferans | I(time^2):recall0.75 | -0.323 | 0.323 | -0.956 | 0.309 | -1.001 | 0.317 |
| Lipaugus vociferans | I(time^2):recall0.9 | -0.064 | 0.306 | -0.665 | 0.536 | -0.21 | 0.834 |
| Lipaugus vociferans | precision0.1 | 8.213 | 0.1 | 8.017 | 8.408 | 82.345 | <.001 |
| Lipaugus vociferans | precision0.1:recall0.1 | -1.087 | 0.327 | -1.728 | -0.447 | -3.328 | <.001 |
| Lipaugus vociferans | precision0.1:recall0.25 | -0.983 | 0.219 | -1.413 | -0.553 | -4.482 | <.001 |
| Lipaugus vociferans | precision0.1:recall0.5 | -0.676 | 0.173 | -1.015 | -0.337 | -3.912 | <.001 |
| Lipaugus vociferans | precision0.1:recall0.75 | -0.366 | 0.153 | -0.667 | -0.066 | -2.392 | 0.017 |
| Lipaugus vociferans | precision0.1:recall0.9 | -0.212 | 0.145 | -0.496 | 0.072 | -1.461 | 0.144 |
| Lipaugus vociferans | precision0.25 | 6.182 | 0.1 | 5.987 | 6.378 | 61.906 | <.001 |
| Lipaugus vociferans | precision0.25:recall0.1 | -0.161 | 0.328 | -0.805 | 0.483 | -0.49 | 0.624 |
| Lipaugus vociferans | precision0.25:recall0.25 | -0.21 | 0.22 | -0.642 | 0.222 | -0.954 | 0.34 |
| Lipaugus vociferans | precision0.25:recall0.5 | -0.063 | 0.173 | -0.403 | 0.277 | -0.361 | 0.718 |
| Lipaugus vociferans | precision0.25:recall0.75 | 0.026 | 0.154 | -0.275 | 0.327 | 0.17 | 0.865 |
| Lipaugus vociferans | precision0.25:recall0.9 | -0.015 | 0.145 | -0.3 | 0.27 | -0.104 | 0.917 |
| Lipaugus vociferans | precision0.5 | 4.831 | 0.101 | 4.633 | 5.03 | 47.648 | <.001 |
| Lipaugus vociferans | precision0.5:recall0.1 | 0.094 | 0.333 | -0.559 | 0.747 | 0.283 | 0.777 |
| Lipaugus vociferans | precision0.5:recall0.25 | -0.046 | 0.224 | -0.485 | 0.393 | -0.206 | 0.837 |
| Lipaugus vociferans | precision0.5:recall0.5 | 0.022 | 0.176 | -0.324 | 0.367 | 0.124 | 0.901 |
| Lipaugus vociferans | precision0.5:recall0.75 | 0.089 | 0.156 | -0.216 | 0.395 | 0.573 | 0.567 |
| Lipaugus vociferans | precision0.5:recall0.9 | 0.009 | 0.147 | -0.28 | 0.298 | 0.06 | 0.952 |
| Lipaugus vociferans | precision0.75 | 3.514 | 0.105 | 3.307 | 3.72 | 33.321 | <.001 |
| Lipaugus vociferans | precision0.75:recall0.1 | 0.072 | 0.347 | -0.607 | 0.751 | 0.207 | 0.836 |
| Lipaugus vociferans | precision0.75:recall0.25 | -0.135 | 0.234 | -0.593 | 0.324 | -0.576 | 0.565 |
| Lipaugus vociferans | precision0.75:recall0.5 | -0.014 | 0.184 | -0.373 | 0.346 | -0.075 | 0.94 |
| Lipaugus vociferans | precision0.75:recall0.75 | 0.023 | 0.162 | -0.295 | 0.342 | 0.144 | 0.885 |
| Lipaugus vociferans | precision0.75:recall0.9 | 0.003 | 0.153 | -0.298 | 0.304 | 0.02 | 0.984 |
| Lipaugus vociferans | precision0.9 | 2.09 | 0.115 | 1.866 | 2.315 | 18.233 | <.001 |
| Lipaugus vociferans | precision0.9:recall0.1 | 0.122 | 0.375 | -0.614 | 0.858 | 0.325 | 0.745 |
| Lipaugus vociferans | precision0.9:recall0.25 | -0.079 | 0.255 | -0.578 | 0.42 | -0.309 | 0.757 |
| Lipaugus vociferans | precision0.9:recall0.5 | -0.022 | 0.2 | -0.413 | 0.37 | -0.109 | 0.913 |
| Lipaugus vociferans | precision0.9:recall0.75 | -0.004 | 0.177 | -0.35 | 0.342 | -0.022 | 0.983 |
| Lipaugus vociferans | precision0.9:recall0.9 | 0.035 | 0.167 | -0.291 | 0.362 | 0.213 | 0.831 |
| Lipaugus vociferans | recall0.1 | -2.365 | 0.325 | -3.002 | -1.727 | -7.266 | <.001 |
| Lipaugus vociferans | recall0.25 | -1.429 | 0.218 | -1.857 | -1.001 | -6.547 | <.001 |
| Lipaugus vociferans | recall0.5 | -0.8 | 0.172 | -1.137 | -0.464 | -4.661 | <.001 |
| Lipaugus vociferans | recall0.75 | -0.395 | 0.152 | -0.693 | -0.097 | -2.6 | 0.009 |
| Lipaugus vociferans | recall0.9 | -0.125 | 0.144 | -0.407 | 0.156 | -0.874 | 0.382 |
| Lipaugus vociferans | time | 10.149 | 0.293 | 9.575 | 10.724 | 34.622 | <.001 |
| Lipaugus vociferans | time:precision0.1 | -10.355 | 0.301 | -10.944 | -9.766 | -34.454 | <.001 |
| Lipaugus vociferans | time:precision0.1:recall0.1 | -0.049 | 0.962 | -1.934 | 1.837 | -0.05 | 0.96 |
| Lipaugus vociferans | time:precision0.1:recall0.25 | -0.08 | 0.651 | -1.355 | 1.195 | -0.123 | 0.902 |
| Lipaugus vociferans | time:precision0.1:recall0.5 | -0.219 | 0.515 | -1.227 | 0.79 | -0.425 | 0.671 |
| Lipaugus vociferans | time:precision0.1:recall0.75 | -0.198 | 0.458 | -1.097 | 0.7 | -0.433 | 0.665 |
| Lipaugus vociferans | time:precision0.1:recall0.9 | 0.069 | 0.435 | -0.785 | 0.922 | 0.158 | 0.875 |
| Lipaugus vociferans | time:precision0.25 | -10.44 | 0.301 | -11.029 | -9.85 | -34.705 | <.001 |
| Lipaugus vociferans | time:precision0.25:recall0.1 | -0.179 | 0.972 | -2.085 | 1.727 | -0.184 | 0.854 |
| Lipaugus vociferans | time:precision0.25:recall0.25 | -0.029 | 0.657 | -1.317 | 1.259 | -0.044 | 0.965 |
| Lipaugus vociferans | time:precision0.25:recall0.5 | -0.208 | 0.518 | -1.224 | 0.808 | -0.401 | 0.689 |
| Lipaugus vociferans | time:precision0.25:recall0.75 | -0.291 | 0.461 | -1.193 | 0.612 | -0.631 | 0.528 |
| Lipaugus vociferans | time:precision0.25:recall0.9 | -0.049 | 0.437 | -0.905 | 0.807 | -0.112 | 0.911 |
| Lipaugus vociferans | time:precision0.5 | -9.956 | 0.308 | -10.56 | -9.352 | -32.305 | <.001 |
| Lipaugus vociferans | time:precision0.5:recall0.1 | -0.451 | 0.996 | -2.404 | 1.502 | -0.452 | 0.651 |
| Lipaugus vociferans | time:precision0.5:recall0.25 | -0.096 | 0.674 | -1.417 | 1.225 | -0.143 | 0.887 |
| Lipaugus vociferans | time:precision0.5:recall0.5 | -0.202 | 0.531 | -1.244 | 0.84 | -0.38 | 0.704 |
| Lipaugus vociferans | time:precision0.5:recall0.75 | -0.32 | 0.472 | -1.245 | 0.605 | -0.678 | 0.497 |
| Lipaugus vociferans | time:precision0.5:recall0.9 | -0.058 | 0.448 | -0.935 | 0.82 | -0.129 | 0.898 |
| Lipaugus vociferans | time:precision0.75 | -8.296 | 0.323 | -8.929 | -7.662 | -25.683 | <.001 |
| Lipaugus vociferans | time:precision0.75:recall0.1 | -0.208 | 1.045 | -2.257 | 1.84 | -0.199 | 0.842 |
| Lipaugus vociferans | time:precision0.75:recall0.25 | 0.408 | 0.71 | -0.982 | 1.799 | 0.576 | 0.565 |
| Lipaugus vociferans | time:precision0.75:recall0.5 | 0.024 | 0.558 | -1.069 | 1.118 | 0.044 | 0.965 |
| Lipaugus vociferans | time:precision0.75:recall0.75 | -0.102 | 0.495 | -1.073 | 0.868 | -0.207 | 0.836 |
| Lipaugus vociferans | time:precision0.75:recall0.9 | -0.027 | 0.469 | -0.947 | 0.892 | -0.058 | 0.954 |
| Lipaugus vociferans | time:precision0.9 | -5.257 | 0.35 | -5.942 | -4.571 | -15.023 | <.001 |
| Lipaugus vociferans | time:precision0.9:recall0.1 | -0.368 | 1.13 | -2.582 | 1.845 | -0.326 | 0.744 |
| Lipaugus vociferans | time:precision0.9:recall0.25 | 0.251 | 0.77 | -1.258 | 1.759 | 0.326 | 0.745 |
| Lipaugus vociferans | time:precision0.9:recall0.5 | 0.062 | 0.605 | -1.125 | 1.248 | 0.102 | 0.919 |
| Lipaugus vociferans | time:precision0.9:recall0.75 | 0.013 | 0.537 | -1.039 | 1.066 | 0.025 | 0.98 |
| Lipaugus vociferans | time:precision0.9:recall0.9 | -0.11 | 0.508 | -1.105 | 0.885 | -0.217 | 0.828 |
| Lipaugus vociferans | time:recall0.1 | 0.196 | 0.953 | -1.672 | 2.063 | 0.205 | 0.837 |
| Lipaugus vociferans | time:recall0.25 | 0.268 | 0.643 | -0.992 | 1.527 | 0.417 | 0.677 |
| Lipaugus vociferans | time:recall0.5 | 0.39 | 0.506 | -0.602 | 1.383 | 0.771 | 0.441 |
| Lipaugus vociferans | time:recall0.75 | 0.344 | 0.45 | -0.537 | 1.225 | 0.766 | 0.444 |
| Lipaugus vociferans | time:recall0.9 | 0.054 | 0.426 | -0.78 | 0.889 | 0.128 | 0.898 |
| Turdus hauxwelli | (Intercept) | -1.572 | 0.015 | -1.602 | -1.542 | -103.552 | <.001 |
| Turdus hauxwelli | I(time^2) | -4.133 | 0.063 | -4.257 | -4.009 | -65.162 | <.001 |
| Turdus hauxwelli | I(time^2):precision0.5 | 2.21 | 0.09 | 2.033 | 2.388 | 24.454 | <.001 |
| Turdus hauxwelli | I(time^2):precision0.5:recall0.1 | 0.043 | 0.219 | -0.386 | 0.472 | 0.196 | 0.845 |
| Turdus hauxwelli | I(time^2):precision0.5:recall0.25 | 0.064 | 0.16 | -0.249 | 0.377 | 0.404 | 0.686 |
| Turdus hauxwelli | I(time^2):precision0.5:recall0.5 | 0.242 | 0.135 | -0.023 | 0.506 | 1.793 | 0.073 |
| Turdus hauxwelli | I(time^2):precision0.5:recall0.75 | 0.317 | 0.128 | 0.067 | 0.567 | 2.484 | 0.013 |
| Turdus hauxwelli | I(time^2):precision0.5:recall0.9 | 0.194 | 0.127 | -0.055 | 0.442 | 1.528 | 0.127 |
| Turdus hauxwelli | I(time^2):precision0.75 | 1.154 | 0.086 | 0.985 | 1.324 | 13.351 | <.001 |
| Turdus hauxwelli | I(time^2):precision0.75:recall0.1 | 0.056 | 0.233 | -0.4 | 0.511 | 0.239 | 0.811 |
| Turdus hauxwelli | I(time^2):precision0.75:recall0.25 | 0.034 | 0.166 | -0.291 | 0.359 | 0.206 | 0.837 |
| Turdus hauxwelli | I(time^2):precision0.75:recall0.5 | 0.091 | 0.137 | -0.177 | 0.359 | 0.667 | 0.505 |
| Turdus hauxwelli | I(time^2):precision0.75:recall0.75 | 0.038 | 0.127 | -0.21 | 0.286 | 0.3 | 0.764 |
| Turdus hauxwelli | I(time^2):precision0.75:recall0.9 | 0.072 | 0.124 | -0.17 | 0.314 | 0.583 | 0.56 |
| Turdus hauxwelli | I(time^2):precision0.9 | 0.49 | 0.088 | 0.318 | 0.663 | 5.564 | <.001 |
| Turdus hauxwelli | I(time^2):precision0.9:recall0.1 | -0.037 | 0.243 | -0.513 | 0.44 | -0.151 | 0.88 |
| Turdus hauxwelli | I(time^2):precision0.9:recall0.25 | -0.043 | 0.172 | -0.381 | 0.295 | -0.25 | 0.802 |
| Turdus hauxwelli | I(time^2):precision0.9:recall0.5 | 0.003 | 0.141 | -0.274 | 0.279 | 0.018 | 0.986 |
| Turdus hauxwelli | I(time^2):precision0.9:recall0.75 | 0.011 | 0.13 | -0.243 | 0.265 | 0.085 | 0.933 |
| Turdus hauxwelli | I(time^2):precision0.9:recall0.9 | -0.006 | 0.126 | -0.253 | 0.242 | -0.045 | 0.964 |
| Turdus hauxwelli | I(time^2):recall0.1 | 1.444 | 0.177 | 1.097 | 1.791 | 8.153 | <.001 |
| Turdus hauxwelli | I(time^2):recall0.25 | 1.155 | 0.125 | 0.91 | 1.4 | 9.229 | <.001 |
| Turdus hauxwelli | I(time^2):recall0.5 | 0.869 | 0.102 | 0.669 | 1.069 | 8.517 | <.001 |
| Turdus hauxwelli | I(time^2):recall0.75 | 0.439 | 0.094 | 0.255 | 0.622 | 4.683 | <.001 |
| Turdus hauxwelli | I(time^2):recall0.9 | 0.188 | 0.091 | 0.01 | 0.366 | 2.065 | 0.039 |
| Turdus hauxwelli | precision0.5 | 2.137 | 0.021 | 2.097 | 2.178 | 103.744 | <.001 |
| Turdus hauxwelli | precision0.5:recall0.1 | -0.824 | 0.053 | -0.927 | -0.721 | -15.679 | <.001 |
| Turdus hauxwelli | precision0.5:recall0.25 | -0.733 | 0.038 | -0.807 | -0.658 | -19.273 | <.001 |
| Turdus hauxwelli | precision0.5:recall0.5 | -0.569 | 0.032 | -0.631 | -0.507 | -17.996 | <.001 |
| Turdus hauxwelli | precision0.5:recall0.75 | -0.317 | 0.03 | -0.375 | -0.259 | -10.727 | <.001 |
| Turdus hauxwelli | precision0.5:recall0.9 | -0.153 | 0.029 | -0.21 | -0.096 | -5.243 | <.001 |
| Turdus hauxwelli | precision0.75 | 0.813 | 0.02 | 0.773 | 0.852 | 40.18 | <.001 |
| Turdus hauxwelli | precision0.75:recall0.1 | -0.188 | 0.056 | -0.299 | -0.078 | -3.336 | <.001 |
| Turdus hauxwelli | precision0.75:recall0.25 | -0.167 | 0.04 | -0.246 | -0.089 | -4.175 | <.001 |
| Turdus hauxwelli | precision0.75:recall0.5 | -0.123 | 0.033 | -0.187 | -0.059 | -3.77 | <.001 |
| Turdus hauxwelli | precision0.75:recall0.75 | -0.075 | 0.03 | -0.134 | -0.017 | -2.517 | 0.012 |
| Turdus hauxwelli | precision0.75:recall0.9 | -0.024 | 0.029 | -0.081 | 0.033 | -0.834 | 0.404 |
| Turdus hauxwelli | precision0.9 | 0.306 | 0.021 | 0.265 | 0.347 | 14.64 | <.001 |
| Turdus hauxwelli | precision0.9:recall0.1 | -0.076 | 0.06 | -0.193 | 0.041 | -1.276 | 0.202 |
| Turdus hauxwelli | precision0.9:recall0.25 | -0.068 | 0.042 | -0.15 | 0.015 | -1.608 | 0.108 |
| Turdus hauxwelli | precision0.9:recall0.5 | -0.045 | 0.034 | -0.112 | 0.022 | -1.327 | 0.185 |
| Turdus hauxwelli | precision0.9:recall0.75 | -0.025 | 0.031 | -0.086 | 0.036 | -0.797 | 0.425 |
| Turdus hauxwelli | precision0.9:recall0.9 | -0.015 | 0.03 | -0.074 | 0.044 | -0.489 | 0.625 |
| Turdus hauxwelli | recall0.1 | -2.386 | 0.044 | -2.472 | -2.3 | -54.414 | <.001 |
| Turdus hauxwelli | recall0.25 | -1.494 | 0.031 | -1.554 | -1.433 | -48.364 | <.001 |
| Turdus hauxwelli | recall0.5 | -0.756 | 0.025 | -0.805 | -0.707 | -30.388 | <.001 |
| Turdus hauxwelli | recall0.75 | -0.319 | 0.023 | -0.363 | -0.274 | -14.089 | <.001 |
| Turdus hauxwelli | recall0.9 | -0.115 | 0.022 | -0.158 | -0.072 | -5.253 | <.001 |
| Turdus hauxwelli | time | 4.77 | 0.067 | 4.639 | 4.901 | 71.358 | <.001 |
| Turdus hauxwelli | time:precision0.5 | -2.576 | 0.094 | -2.761 | -2.392 | -27.401 | <.001 |
| Turdus hauxwelli | time:precision0.5:recall0.1 | -0.041 | 0.23 | -0.492 | 0.411 | -0.177 | 0.86 |
| Turdus hauxwelli | time:precision0.5:recall0.25 | -0.099 | 0.168 | -0.428 | 0.23 | -0.59 | 0.555 |
| Turdus hauxwelli | time:precision0.5:recall0.5 | -0.251 | 0.141 | -0.527 | 0.026 | -1.774 | 0.076 |
| Turdus hauxwelli | time:precision0.5:recall0.75 | -0.359 | 0.133 | -0.62 | -0.098 | -2.698 | 0.007 |
| Turdus hauxwelli | time:precision0.5:recall0.9 | -0.202 | 0.132 | -0.461 | 0.057 | -1.527 | 0.127 |
| Turdus hauxwelli | time:precision0.75 | -1.355 | 0.091 | -1.532 | -1.177 | -14.959 | <.001 |
| Turdus hauxwelli | time:precision0.75:recall0.1 | -0.066 | 0.245 | -0.547 | 0.414 | -0.27 | 0.787 |
| Turdus hauxwelli | time:precision0.75:recall0.25 | -0.048 | 0.175 | -0.391 | 0.295 | -0.275 | 0.783 |
| Turdus hauxwelli | time:precision0.75:recall0.5 | -0.089 | 0.144 | -0.371 | 0.192 | -0.622 | 0.534 |
| Turdus hauxwelli | time:precision0.75:recall0.75 | -0.035 | 0.133 | -0.295 | 0.225 | -0.263 | 0.793 |
| Turdus hauxwelli | time:precision0.75:recall0.9 | -0.071 | 0.129 | -0.324 | 0.183 | -0.547 | 0.584 |
| Turdus hauxwelli | time:precision0.9 | -0.577 | 0.093 | -0.759 | -0.396 | -6.235 | <.001 |
| Turdus hauxwelli | time:precision0.9:recall0.1 | 0.05 | 0.257 | -0.455 | 0.554 | 0.193 | 0.847 |
| Turdus hauxwelli | time:precision0.9:recall0.25 | 0.052 | 0.182 | -0.305 | 0.41 | 0.286 | 0.775 |
| Turdus hauxwelli | time:precision0.9:recall0.5 | 0.005 | 0.149 | -0.287 | 0.297 | 0.034 | 0.973 |
| Turdus hauxwelli | time:precision0.9:recall0.75 | -0.005 | 0.137 | -0.273 | 0.263 | -0.038 | 0.97 |
| Turdus hauxwelli | time:precision0.9:recall0.9 | 0.012 | 0.133 | -0.248 | 0.272 | 0.091 | 0.927 |
| Turdus hauxwelli | time:recall0.1 | -1.644 | 0.188 | -2.013 | -1.276 | -8.752 | <.001 |
| Turdus hauxwelli | time:recall0.25 | -1.292 | 0.133 | -1.552 | -1.031 | -9.727 | <.001 |
| Turdus hauxwelli | time:recall0.5 | -0.975 | 0.108 | -1.187 | -0.763 | -9.03 | <.001 |
| Turdus hauxwelli | time:recall0.75 | -0.51 | 0.099 | -0.703 | -0.316 | -5.153 | <.001 |
| Turdus hauxwelli | time:recall0.9 | -0.223 | 0.096 | -0.411 | -0.035 | -2.325 | 0.02 |
